# Ablating Ku70 phosphorylation results in defective DNA damage repair and spontaneous induction of hepatocellular carcinoma

**DOI:** 10.1101/2021.03.03.433432

**Authors:** Janapriya Saha, Jinsung Bae, Shih-Ya Wang, Lori J. Chappell, Purva Gopal, Anthony J. Davis

**Affiliations:** Division of Molecular Radiation Biology, Department of Radiation Oncology, UT Southwestern Medical Center, Dallas, TX; KBR, Houston, TX; Department of Pathology, UT Southwestern Medical Center, Dallas, TX

**Keywords:** Ku70, Hepatocellular carcinoma, DNA damage, Chromosomal instability, radiation

## Abstract

Multiple pathways mediate the repair of DNA double-strand break (DSB), with numerous mechanisms responsible for driving choice between the pathways. Previously, we reported that phosphorylation of the non-homologous end joining (NHEJ) factor, Ku70, is required for the dissociation of the Ku heterodimer from DNA ends to allow DSB repair via homologous recombination (HR). A knock-in mouse, in which phosphorylation is ablated in the three conserved sites of Ku70 (Ku70^3A/3A^), was generated in order to test the hypothesis that Ku70 phosphorylation is required for initiation of HR and that blocking this process results in enhanced genomic instability and tumorigenesis. Here, we show that Ku70^3A/3A^ mice develop spontaneous and have accelerated chemical-induced hepatocellular carcinoma (HCC) compared to wild-type (Ku70^+/+^) littermates. The HCC tumors from the Ku70^3A/3A^ mice have increased γH2AX and 8-oxo-G staining, suggesting DNA repair is decreased in these mice. Spontaneous transformed cell lines from Ku70^3A/3A^ mice are more radiosensitive, have a significant decrease in DNA end resection, and are more sensitive to the DNA cross-linking agent mitomycin C compared to cells from Ku70^+/+^ littermates. Collectively, these findings demonstrate that phosphorylation-mediated dissociation of Ku heterodimer from DNA ends is required for efficient DNA damage repair and disruption of this process results in genomic instability and accelerated development of HCC.

## INTRODUCTION

The Ku protein, a heterodimer consisting of the subunits Ku70 and Ku80, plays critical roles in cellular processes such as non-homologous end joining (NHEJ), V(D)J recombination, apoptosis, telomere maintenance, aging, senescence, and DNA replication (Davis et al., 2014; Doherty and Jackson, 2001; Fell and Schild-Poulter, 2015; Frit et al., 2019). Ku is an abundant, highly conserved protein that possesses extremely high affinity for double-stranded (ds) DNA ends, sliding onto DNA via its ring-like structure that is generated by the intertwining of the subunits. It can bind various dsDNA ends, including those with blunt, overhangs, and hairpins, and heterogeneous ends generated by ionizing radiation (IR). The most defined function of Ku is its role in the maintenance of genome integrity via initiation of the NHEJ pathway, the main DNA double strand break (DSB) repair pathway in mammalian cells (Davis et al., 2014; Davis and Chen, 2013). Following induction of a DSB, Ku70/80 binds to the DNA damage site within seconds of its creation and does so in all cell cycle phases (Davis et al., 2014; Mari et al., 2006; Shao et al., 2012). Ku recruits and activates the DNA-PK_cs_ kinase to the damage site, where the DNA-PK complex (DNA-PK_cs_-Ku-DNA) initiates the DNA damage response and chromatin remodeling (Davis et al., 2014; Davis and Chen, 2013; Lu et al., 2019). Once bound to the DSB, Ku performs its primary function in NHEJ, which is to serve as a scaffold to recruit the NHEJ machinery to the DNA lesion (Davis and Chen, 2013).

Delineating Ku’s functions has been difficult in human cells because depletion of Ku70 is lethal and induced loss of Ku80 causes rapid cell death accompanied by massive telomere loss (Fattah et al., 2008; Wang et al., 2009). However, Ku-deficient mouse models have been instrumental in examining its function in multiple pathways. Ku70^-/-^, Ku80^-/-^, Ku70^-/-^Ku80^-/-^ mice are viable and reproduce, but they show profound premature aging and are 40-60% the size of wild-type littermates (Gu et al., 1997b; Li et al., 1998; Li et al., 2007; Nussenzweig et al., 1996; Ouyang et al., 1997; Vogel et al., 1999; Zhu et al., 1996). Consistent with the aging and growth defect phenotypes, mouse embryonic fibroblasts (MEFs) derived from Ku70^-/-^ and Ku80^-/-^ mice display premature senescence which is associated with an early loss of proliferating cells, a prolonged doubling time, intact cell cycle checkpoints in response to DNA damage, and an accumulation of nondividing cells (Gu et al., 1997b; Li et al., 1998; Nussenzweig et al., 1996). Ku80^-/-^ mice also exhibit premature age-specific changes including hepatocellular degeneration, hepatocellular inclusions, and hepatic hyperplastic foci (Vogel et al., 1999). The role of Ku protecting the genome by directing DSB repair via NHEJ has also been supported by studies in various Ku-deficient rodent cell lines. Ku70 and Ku80 deficient MEFs and Chinese hamster ovary (CHO) cells are sensitive to DNA damage generated by IR (Gu et al., 1997a; Lee et al., 2016; Li et al., 1998; Nussenzweig et al., 1996; Taccioli et al., 1994; Vogel et al., 1999). V(D)J recombination requires NHEJ and not surprisingly, Ku-deficient mice are defective in this process (Gu et al., 1997b; Nussenzweig et al., 1996; Ouyang et al., 1997; Zhu et al., 1996). T and B lymphocyte development is arrested at early progenitor stages and there is a profound deficiency in V(D)J rearrangement in Ku80^-/-^ mice (Nussenzweig et al., 1996; Zhu et al., 1996). Ku70^-/-^ mice lack mature B cells or serum immunoglobulin but, develop small populations of thymic cells (Gu et al., 1997b; Li et al., 1998).

Although it is firmly established that Ku plays a pivotal role in maintaining genome integrity, its role as a tumor suppressor is not clearly defined. Ku70^-/-^ mice were found to have a significant incidence of thymic lymphomas (Gu et al., 1997b; Li et al., 1998). However, a subsequent study with Ku70^-/-^, Ku80^-/-^, and Ku70^-/-^Ku80^-/-^ mice with the same genetic background and raised in a similar environment found early aging in each cohort of mice, but without substantially increased cancer levels (Li et al., 2007). Ku80^−/−^combined with p53^−/−^ results in early incidence of pro-B-cell lymphoma that resembles Burkitt’s lymphoma (Difilippantonio et al., 2000). Haplo-insufficiency of Ku80 in PARP^-/-^ mice (Ku80^+/-^PARP^-/-^ mice) promotes the development of hepatocellular adenoma and hepatocellular carcinoma (HCC), which correlated with elevated chromosomal aberrations (Tong et al., 2002). Ku70^-/-^ mice had accelerated HCC development compared to wild-type and Ku70^+/-^ littermates after treatment with the liver carcinogen diethylnitrasamine (DEN) (Teoh et al., 2008). Cytogenetic studies found that Ku70^-/-^ HCCs harbored clonal increases in numerical and structural aberrations of chromosomes, including fragments, end-to-end fusions, and recurrent nonreciprocal translocations. A deficiency of toll-like receptor 4 (TLR4)-mediated immune activities resulted in increased DEN-induced HCC development and these tumors showed increased DNA damage, which was due to a decrease in TLR-mediated expression of Ku70 and Ku80 (Wang et al., 2013). The role of Ku70/80 in promoting tumor suppression in humans is mostly unknown. No human patient has yet been identified with a verified disease-causing mutation or loss of Ku, but a number of reports indicate that single nucleotide polymorphisms in Ku70 or Ku80 potentially contribute to different types of cancer including HCC, breast, lung, oral, bladder, renal, and colorectal cancers (reviewed in (Sishc and Davis, 2017)).

A secondary function that Ku performs at DSBs is a general role of binding to and maintaining the stability of the broken DNA ends in order to protect them from non-specific processing, which if left unchecked could potentially lead to loss of genomic sequences resulting in the generation of chromosomal aberrations and genomic instability (Davis et al., 2014; Shao et al., 2012). For example, Ku blocks DNA end processing enzymes including exonuclease 1 and the Mre11/Rad50/Nbs1 complex *in vitro* (Sun et al., 2012). Ku maintains the two ends of the broken DNA molecule together via forming a synaptic complex *in vitro* and is also required for the positional stability of DSB ends *in vivo* (Cary et al., 1997; DeFazio et al., 2002; Soutoglou et al., 2007). Ku binds to DSBs in all cell cycle phases including in S phase, the cell cycle stage where homologous recombination (HR) is the preferred DSB repair pathway (Lee et al., 2016; Shao et al., 2012). Previously, we aimed to delineate the mechanism that drives Ku’s dissociation from DSBs in order to allow HR to occur and we identified that phosphorylation of Ku70 mediates this process (Lee et al., 2016). We found that blocking Ku70 phosphorylation led to a significant decrease in DNA end resection and HR in S phase of the cell cycle (Lee et al., 2016). These results collectively drive our proposed model that phosphorylation-mediated dissociation of Ku from DSBs is one of the mechanisms that modulates DSB repair pathway choice in mammalian cells. However, the importance and physiological function of Ku70 phosphorylation *in vivo* is unknown. Using a mouse model with knock-in alanine substitutions (Ku70^3A/3A^ and Ku70^3A/+^) of the conserved Ku70 phosphorylation sites, we aimed to test our hypothesis that initiation of HR requires dissociation of Ku70/80 from DSB ends and that blocking this process results in enhanced genomic instability and tumorigenesis. Here, we report findings that blocking Ku70 phosphorylation results in increased incidence of spontaneous and accelerated induction of chemical-induced HCC. Moreover, we found that Ku70^3A/3A^ mice and MEFs are radiosensitive and that Ku70^3A/3A^ MEFs have decreased DNA end resection and increased sensitivity to the DNA cross-linking agent mitomycin C (MMC). Collectively, the results suggest that blocking Ku70 phosphorylation results in defective DNA repair and that this drives genomic instability and development of HCC.

## MATERIALS AND METHODS

### Cell culture

Primary MEFs isolated from E13.5 embryos from Ku70^+/+^ and Ku70^3A/3A^ mice were cultured in Hyclone α-MEM (GE Life Sciences) supplemented with 10% fetal bovine serum (Corning) and 1x penicillin/streptomycin (GE Life Sciences) and initially grown in an atmosphere of 3% O_2_ and 10% CO_2_ at 37°C in order to drive spontaneous transformation. Following transformation, the primary MEFs were grown in the same media in an atmosphere of 5% CO_2_. Ku70^-/-^ MEFs (Li et al., 1998) were grown in Hyclone α-MEM supplemented with 5% fetal bovine serum, 5% newborn calf serum (GE Life Sciences), and 1x penicillin/streptomycin in an atmosphere of 5% CO_2_ at 37°C. Ku70^-/-^ MEFs complemented with mouse wild-type Ku70 and Ku70 3A were maintained in the same media but supplemented with 10 µg/ml hygromycin (Corning).

### Irradiation

Cells were irradiated with γ-rays generated by a Mark 1 ^137^Cs irradiator (JL Shepherd and Associates) at the doses denoted in the figures.

### Mouse Ku70 mutagenesis and generation of stable cell lines

Mouse Ku70 cDNA was PCR amplified using the primers mKu70-Bam-ATG: 5’-GGCGGATCCATGTCAGAGTGGGAGTCCTAC-3’ and mKu70-Stop-Xho: 5’-GGCCTCGAGTCAGTTCTTCTCCAAGTGTCTG-3’ and then subcloned into the pCDNA3.1-YFP-F2 mammalian expression vector using BamHI and XhoI following standard protocols. PCR-directed mutagenesis using complimentary oligonucleotides was then used to generate mouse Ku70 3A. The following primers were used for the mutagenesis: mKu70-307A: 5’-GACTTTTAATGTAAACGCCGGCAGTCTACTCC-3’ and mKu70-314A/316A: 5’-CAGTCTACTCCTGCCTGCTGACGCCAAGCGGTCTCTGAC-3’ with the mKu70-307A primer set used first and then the mKu70-314A/316A primer set. To make Ku70^-/-^ MEFs that stably express wild-type and Ku70 3A, 4 μg of linearized plasmid containing wild-type or Ku70 3A cDNA was transfected using Amaxa nucleofector solution V (Lonza) and program T-020. Cells stably expressing mKu70 were selected using 800 μg/mL hygromycin.

### RPA/Rad51 foci kinetics

IR-induced RPA/Rad51 kinetics were determined as previously outlined (Saha et al., 2017). Cells were grown on coverslips one day prior to the experiment. On the day of the experiment, EdU (Molecular Probes) was added to a final concentration of 30 μM and incubated for 40 min under the normal growth conditions. After 40 min, the cells were washed twice with 1x PBS and fresh media was added. The cells were then subjected to 8 Gy of γ-rays. The cells were allowed to recover for different time intervals, as denoted in the figures. The cells were washed twice with ice cold 1x PBS and then treated with freshly made CSK extraction buffer (10 mM HEPES pH 7.4, 300 mM Sucrose, 100 mM NaCl, 3 mM MgCl_2,_ 0.1% Triton) for 10 min on ice. The cells were then washed 5 times with ice cold 1x PBS. The cells were fixed with 4% paraformaldehyde (PFA) (in 1x PBS) for 20 min at ambient temperature, then washed 5 times with 1x PBS at 5-minute intervals, followed by incubation in 0.5% Triton X-100 in ice-cold PBS on ice for 10 min. The cells were washed 5 times with 1x PBS and incubated in blocking solution (5% goat serum (Jackson Immuno Research) in 1x PBS) overnight. RPA and Rad51 foci were detected using antibodies against RPA34 (NA-19L, EMD Millipore) or Rad51 (SC-8349, Santa Cruz), respectively, in 5% goat serum in 1x PBS. Following a 2 hr incubation, the cells were washed 5 times with wash buffer (1% BSA in 1x PBS), with each wash lasting 5 minutes. EdU detection was performed using the Click-iT reaction cocktail according to the manufacturer’s protocol (Molecular Probes). After incubation with the Click-iT reaction cocktail in the dark for 1 hr, the cells were washed 5 times with wash buffer. The cells were then incubated with Alexa Fluor 488 (1:1000) (Molecular Probes) secondary antibody in 1% BSA, 2.5% goat serum in 1x PBS for 1 hr in the dark, followed by 5 washes with wash buffer. After the last wash, the cells were mounted in VectaShield mounting medium containing DAPI. The images were acquired using a Zeiss Axio Imager fluorescence microscope utilizing a 63x oil objective.

### Mice

All mice were approved by and handled in accordance with the guidelines of the Institutional Animal Care and Use Committee (IACUC) at UT Southwestern Medical center (UTSW). All mice were bred and maintained in a specific pathogen free SPF barrier vivarium at UTSW. All experiments were conducted with age and sex matched mice and treatment groups were allocated randomly.

### Generation of Ku70^3A/3A^ knock-in mice

ES cell targeting and generation of chimeric mice were performed at the transgenic core facility at UTSW. Specifically, C57BL/6N females (Charles River) were induced to superovulate by injecting pregnant mare serum gonadotropin (PMSG) and human chorionic gonadotropin (HCG) and then mated with C57BL/6N males to collect the zygotes for microinjection. A mixtures of Cas9-mRNA (Sigma)(50ng/μL), synthesized gRNA (Sigma) (25ng/μL) and template single-strand oligodeoxynucleotide (ssODN) (Sigma)(25ng/μL) were injected into pronuclear of one-cell zygotes. The Ku70 gRNA sequence utilized is 5’-ACACCAAGCGGTCTCTGGTAGG-3’ and the template ssODN (donor oligo) sequence is 5’-GAACCAGTGAAAACCAAGACAAGGACTTTTAATGTAAAC<u>G</u>CCGGCAGTCTACTCCTGCCT <u>GC</u>TGAC<u>G</u>CCAAGCGGTCTCTGGTAGGTGGCTAACCTTTCCTACCGAATCTTGTTTAAGA-3’ (the mutation sites are underlined). After microinjection, the zygotes were implanted into ICR recipients, which carried them to term. Seven chimeras were obtained and then backcrossed twice with wild-type C57BL/6J mice (Jackson Laboratory) in order to limit potential off-target mutations generated by CRISPR/Cas9. For sequencing and PCR genotyping, DNA isolated from either the mouse tail or ear was used. Specifically, a small piece of tail or ear (approximately 2 mm) was excised and placed and in an Eppendorf tube. 400 µL of tail buffer (100 mM Tris (pH 8.0), 5 mM EDTA, 2% SDS, and 20 mM NaCl) and 6 µL Proteinase K (10 mg/mL) was added to each tube and incubated at 55°C with shaking overnight or in a 55°C block until the tail/ear was dissolved. The tubes were then centrifuged at 13,600 rpm for 30 minutes, and the supernatant transferred to a new Eppendorf tube. 400 µL isopropanol was added to each tube, the samples were mixed via shaking 50 times, then centrifuged at 13,600 rpm for 5 minutes, and the supernatant was discarded. The samples were then air-dried in a chemical hood at ambient temperature. 100 µL of 1x TE buffer was added to each sample, incubated at 55°C until the DNA was dissolved, and the DNA concentrations were determined. For sequencing, the following primers were used to amplify Ku70 sequencing, mKu70-1129F (5’-TTGGACAGAGTAACGCAGACTGG-3’) and mKu70-11802R (5’-CTCGTGCATGCCTCATGCATGC-3’). Sequencing was performed at the UTSW Sanger Sequencing Core. For genotyping of mice via PCR analysis, the following primer sets were used: Wild-type-primers mKu70-WT (5’-CGGCAGTCTACTCCTGCCTAG-3’) and mKu70-11802R and Ku70-3A-primers mKu70-314A (5’-CGGCAGTCTACTCCTGCCTGC-3’).

### Monitoring of mice and examination of tumorigenesis

Ku70^3A/3A^, Ku70^3A/+^, and Ku70^+/+^ littermates were maintained and monitored for up to 28 months. Their weight and activity were monitored monthly until one-year of age and bi-weekly thereafter. Necropsy was performed at various time points and the mice were examined for gross tumors. All tissues with tumors were collected and fixed with 4% neutralized PFA for 24 hours. Clinical histopathological of H&E-stained sections were performed blindly. A total of 42 Ku70^3A/3A^ mice (22 males and 20 females), 72 Ku70^3A/+^ mice (44 males and 28 females), and 30 Ku70^+/+^ wild-type mice (21 males and 9 females) were included in the final analysis.

### DEN-induced HCC model

Male C57BL/6 Ku70^3A/3A^ and Ku70^+/+^ littermates were injected with DEN (Sigma) (25 mg/kg i.p.) intraperitoneally (i.p.) at day 14 postpartum. All mice were monitored weekly for weight and activity and sacked at different intervals (4, 6, 9 months) as appropriate to test for liver disease. 9 months following DEN injection, all surviving mice from all genotypes were euthanized with CO_2_ and their liver tissue was collected and fixed. DEN-induced liver tumor numbers were determined visually, and size determined using a Vernier caliper.

### Immunohistochemistry (IHC)

The fixed tissues samples were processed for paraffin sectioning and stained with hematoxylin-eosin (H&E) according to standard protocol by the Histo-Pathology Core at UTSW. IHC staining was performed using the following antibodies, Phospho-Histone H2A.X (Ser139) (20E3) Rabbit mAb (Cell Signaling), Ki-67 (D3B5) Rabbit mAb (Cell Signaling), and DNA/RNA damage antibody (8-oxo-G) [15A3] (Abcam) by the Histo-Pathology Core at UTSW.

### Total body irradiation survival experiments

Healthy 8-week-old Ku70^3A/3A^ mice and Ku70^+/+^ littermates were irradiated with 9 Gy total body irradiation (TBI) utilizing the self-contained X-ray irradiation system (XRAD 320, Precision X-Ray) with a maximum output of 320 kV. Their weight and activity were monitored daily until 30 days post-irradiation. The mice were sacked when they lost more than 20% of their initial body weight, their Body Conditioning Score (BCS) fell below 2%, or at the date of experimental termination. P-values for survival curves were determined from the Kaplan–Meier survival curves by use of the log-rank test. The number (N) for each group and batch number of experiments are indicated in the corresponding figure legends. Survival curves and histogram figures were generated by GraphPad Prism software (GraphPAD Software, San Diego).

### Clonogenic survival assays

For clonogenic survival assays, DC-1 (Ku70 null) and DC-1 complemented with wild-type mouse Ku70 or Ku70 3A protein or spontaneously transformed MEFs from Ku70^+/+^ or Ku70^3A/3A^ (clones #51 and #57) were treated with increasing doses of DNA damaging agents (γ-rays or MMC, see figures for doses) and then plated on 60-mm plastic Petri dishes (Greiner Bio-One). After 10 days, the cells were stained with 0.1% crystal violet in a 100% methanol solution. Colonies were scored and the mean value for triplicate culture dishes was determined. Cell survival was normalized to plating efficiency of untreated controls for each cell type.

### Chromosomal abnormalities

For chromosomal abnormalities, spontaneously transformed primary MEFs (passages 8-10) were incubated with 200 ng/mL MMC (Sigma) for 20 hr followed by 0.1 μg/mL Demecolcine Solution (Sigma) in the last 4 hr. After the treatment, the cell media was collected. The cells were then washed once with 1x PBS and trypsinized with 1x 0.25% Trypsin-EDTA (Gibco). The trypsin was neutralized using the media collected before in a 15ml Falcon tube. Cells were then centrifuged at 1,000 rpm for 5 min at RT. The supernatant was discarded and the pellet was resuspended in 10 mL of prewarmed (37^○^C) 75 mM KCl solution by gently tapping and incubated in a heated water bath (37^○^C) for 15 min. 1ml of freshly prepared and pre-cooled fixative (methanol:acetic acid 3:1) was added to the tube and the solution was mixed gently. The samples were centrifuged at 1,000 rpm for 5 min at 4°C and the supernatants were discarded. The pre-cooled fixative was slowly added to resuspend the cell pellet with gentle tapping until the total volume was 10 ml. The tubes were stored at −20^○^C until further processing. On the day of processing, cells were centrifuged at 1,000 rpm for 5 min at 4°C. The supernatant was aspirated, and a small volume of buffer was left to attain a suitable cell density on the slide (2-5 x 10^5^ cells/mL). The cell suspension was then dropped onto pre-cleaned slides (Gold Seal) using a plastic pipette from about 1-meter height above the slide and the slides were allowed to air dry overnight. The rest of the cell suspension was stored in 5 ml of fixative solution at −20^○^C. The slides were then stained with 5% Giemsa solution (Gurr Buffer: KH_2_PO_4_ (0.54 g/L) and Na_2_HPO_4_ (0.57 g/L). 2 volumes of KH_2_PO_4_ and 1 volume of Na_2_HPO_4_ were mixed to make a solution that was 46.5 mL, and to this 2.5 mL Giemsa stain (Gibco) and 1 mL of acetone was added and mixed together. The staining solution was placed in a coplin jar and the slides were added to the jar to stain for 5 min at ambient temperature. The slides were then washed with running water and allowed to air dry. For mounting, a drop of Permount (Fisher Scientific) was added to the slide and a cover glass placed on top. The chromosomes were analyzed and scored with a Zeiss Axio Imager fluorescence microscope using a 100x oil immersion lens.

### Isolation of splenic immune cells

Total immune cells from the spleen were isolated following previously reported methods (Zhang et al., 2017). Specifically, the spleen was rinsed with 5 ml of ice-cold PBS and placed into a petri dish with 10 mL PBS + 0.1% BSA. Single-cell suspension was created by forcing the spleen through a stainless wire screen with a syringe plunger. Cell debris was removed by passing the cell suspension through a BD Falcon (BD Biosciences, Franklin Lakes, NJ, USA) 100-mL cell strainer to a 50-mL conical tube. The cells were washed with 50 mL of ice-cold PBS + BSA and then pelleted by centrifugation. The cell pellet was resuspended in 5 mL (1X) red blood cell lysis buffer (eBioscience). After 5 min at ambient temperature, 25 mL cold HBSS +1% FBS was added to the 50-mL tube to stop lysis. The cell suspension was filtered with a 70 μm cell strainer. The cell number and viability were determined using Beckman Coulter Z2 Coulter® Particle Count and Size Analyzer. The cells were then centrifuged at 350 x g for 5 min at 4°C. The supernatant was discarded, and the cells were resuspended with the right amount of HBSS +1% FBS to reach final concentration of 1 x 10^8^ cells/mL for staining.

### Staining of splenic immune cells

For cell surface staining, lymphocytes isolated from spleens were mixed with anti-mouse CD16/32 mAbs on ice for 5 min to block Fc receptors. Then, cells were mixed with Ab cocktail containing mAbs against specific cell-surface markers and incubated on ice for 30 min in the dark. The following antibodies were used (all purchased from eBioscience (San Diego, CA, USA, now Affymetrix, part of Thermo Fisher Scientific), Panel 1 (Ly6G-FITC, F4/80-PE, CD11b-PE-Cy7, Ly6C-PerCP-Cy5.5, and CD11c-APC) and Panel 2 (CD19-FITC, CD25-PE, CD3-PE-Cy7, CD8-PerCP-Cy5.5, CD4-AF700, and CD49b-eFluor450). The stained samples were transferred to 5-mL flow tubes and 200 µL of staining buffer was added to each sample. The samples were run on a FACS Calibur (BD) and data analyzed using Flo Jo software by the Flow Cytometry Core at UTSW. Compensation was performed on the FACS Calibur (BD) flow cytometer at the beginning of each experiment.

## Results

### Three of five Ku70 phosphorylation sites are found in mice and their function is conserved

Previously, we reported that phosphorylation at a cluster of sites in the junction of the pillar and bridge regions of human Ku70 is required for the dissociation of the Ku heterodimer from DSBs (Lee et al., 2016). Blocking phosphorylation of human Ku70 leads to sustained retention of Ku at DSBs, resulting in a significant decrease in DNA end resection and HR in S phase of the cell cycle (Lee et al., 2016). An alignment of DNA sequences of Ku70 orthologues found that the five sites are conserved only in primates, whereas T307, S314, and T316 are conserved in vertebrates, including in mice (Fig 1A **and Sup. Fig. 1**). To test the functionality of the conserved sites in mice, the three sites were mutated in mouse Ku70 cDNA to alanine to ablate phosphorylation and then stably expressed in the mouse Ku70^-/-^ cell line DC-1. Similar to our previous data with the human Ku70 5A mutant, the mouse Ku70 3A protein is retained at micro-irradiation generated DSBs (Fig. 1B) and caused moderate radiosensitivity compared to DC-1 cells complemented with wild-type mouse Ku70 (Fig. 1C). Furthermore, cells expressing mouse Ku70 3A showed decreased DNA end resection and ongoing HR, as monitored by IR-induced RPA and Rad51 foci resolution, respectively (Fig. 1D **and** E, **Sup. Fig. 2A and B**). Collectively, the data show that three of the five Ku70 phosphorylation sites are conserved in mice and show similar functions as the human sites.

**Figure 1.**
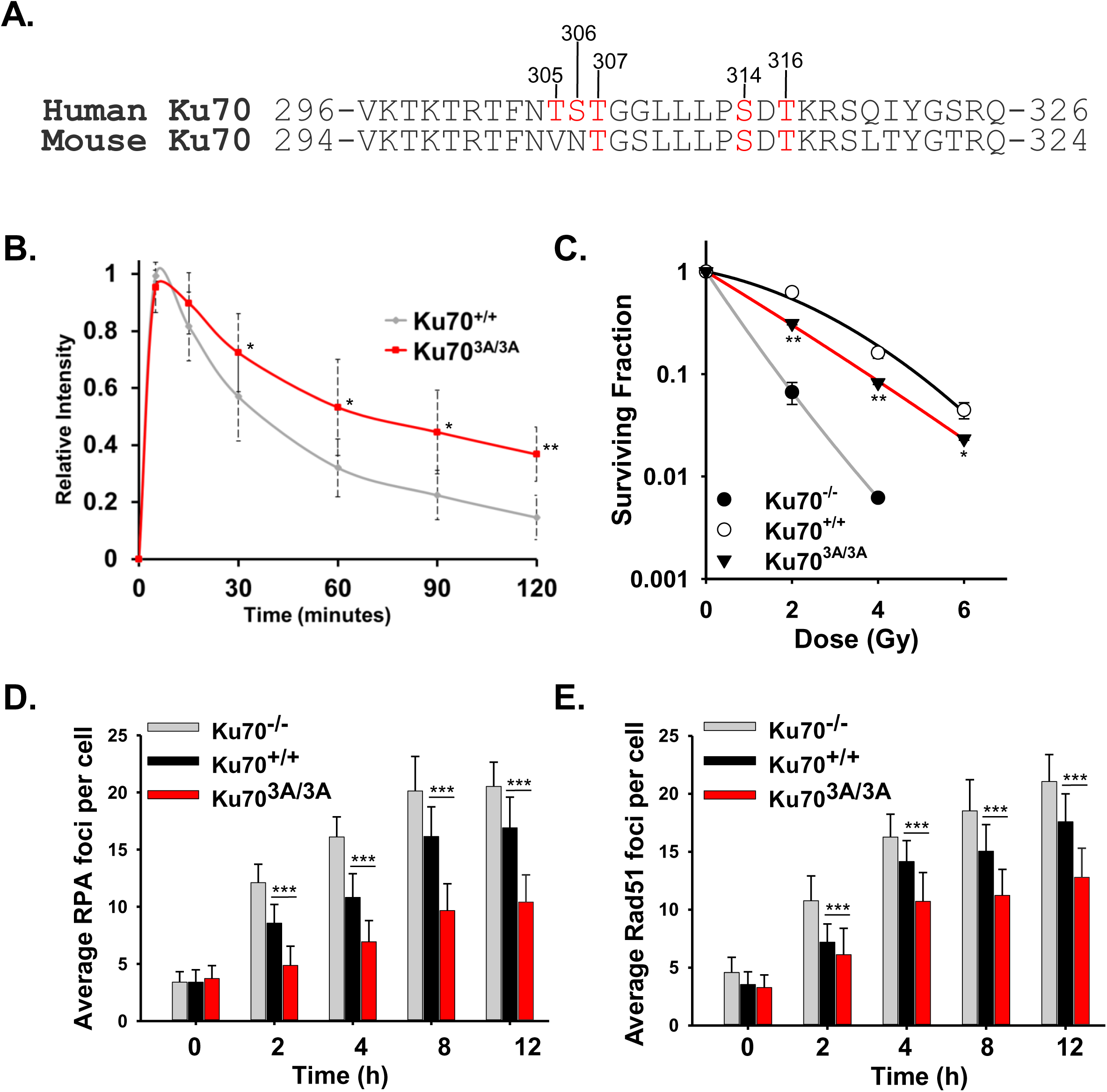
Three of the five Ku70 phosphorylation sites are conserved in mice and are functional. **A.** Alignment of human and mouse Ku70 DNA sequences. Phosphorylation sites are in red. **B.** Relative fluorescence intensity of YFP-tagged Ku70 and Ku70 3A proteins in Ku70^-/-^ MEFs at DSBs after micro-irradiation. Student’s t-test was performed to asses statistical significance (*p < 0.05 and **p < 0.01). **C.** Clonogenic survival assays were performed to compare the radiation sensitivities of Ku70^-/-^ MEFs or Ku70^-/-^ MEFs complemented with Ku70 wild-type (Ku70^+/+^) or 3A (Ku70^3A/3A^). Cells were irradiated at the indicated doses and plated for analysis of survival and colony-forming ability. Error bars denote SD values and Student’s t-test was performed to assess statistical significance (*p < 0.05 and **p < 0.01). (**D and E**) Immunostaining of Ku70^−/−^ MEFs or Ku70^−/−^ MEFs complemented with Ku70 wild-type (Ku70^+/+^) or 3A (Ku70^3A/3A^) after 8 Gy. Cells were pre-extracted and fixed 2, 4, 8, or 12 h after IR and immunostained for RPA (**D**) or Rad51 (**E**) in EdU positive cells (S phase). RPA and Rad51 foci were counted for each cell and averaged. Student’s t-test was performed to assess statistical significance (***p < 0.001). Error bars denote standard error of the mean (SEM) for at least 3 independent experiments.

### Generation of Ku70^3A/3A^ mice

To examine the impact of Ku70 phosphorylation *in vivo*, we generated a C57BL/6 mouse model with alanine substitutions in the three conserved phosphorylation sites (Ku70^3A/3A^) using CRISPR/Cas9 (Fig. 2A-D). Specifically, a mixture of Cas9-mRNA, synthesized gRNA for Ku70, and template ssODN with the sequence for Ku70 with the appropriate mutations were injected into one-cell zygotes of C57BL/6 mice in the pronuclear stage. After microinjection, the zygotes were implanted into ICR recipients, which carried them to term. Seven chimeras were obtained and then backcrossed twice with wild-type C57BL/6 mice in order to limit potential off-target mutations generated by CRISPR/Cas9. The resulting Ku70^3A/+^ mice were used and intercrosses between Ku70^3A/+^ mice produced Ku70^3A/3A^ mice at the expected Mendelian ratio. Furthermore, the Ku70^3A/3A^ male and female mice were fertile. Sequencing and PCR analysis validated the Ku70^3A/+^ and Ku70^3A/3A^ genotypes and further studies were performed using three separate founder lines (Fig. 2B **and** C). Ku70^+/+^, Ku70^3A/+^, Ku70^3A/3A^ mice were tracked for growth once a month until 4 months of age and then every 2 months until the mice were 24+ months old. All three genotypes (males and females) developed normally (Fig. 2D **and** E **and Sup. Fig. 3**), and life span analysis showed that there was no significant difference between the three groups of mice up to 24 months of age (Fig. 2F). Given that the severe growth retardation and premature aging phenotype of Ku70-deficient mice (Gu et al., 1997a; Gu et al., 1997b; Li et al., 1998; Li et al., 2007), these data suggest that knock-in mutations at the three phosphorylation sites on Ku70 does not ablate Ku’s role in telomere maintenance.

**Figure 2.**
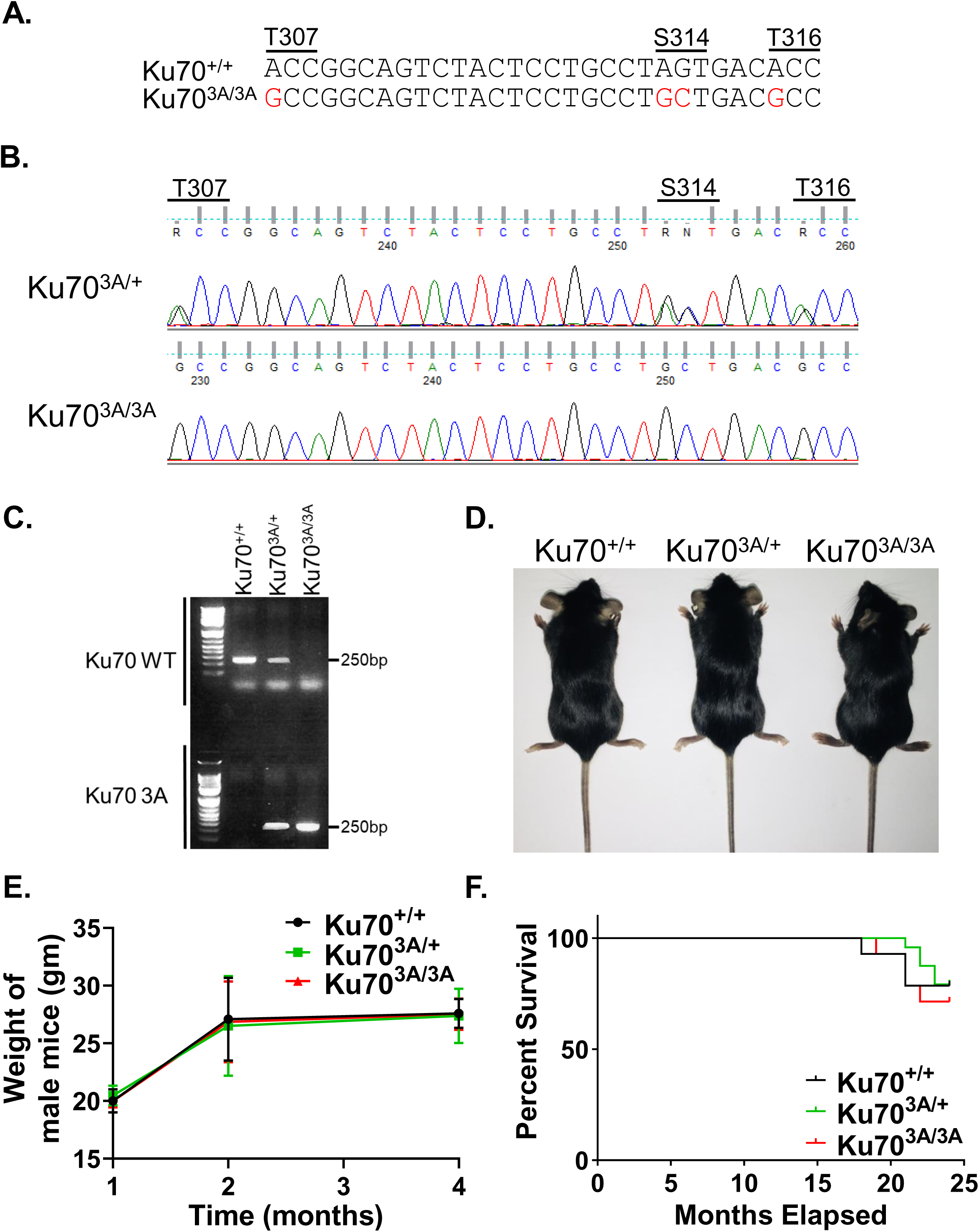
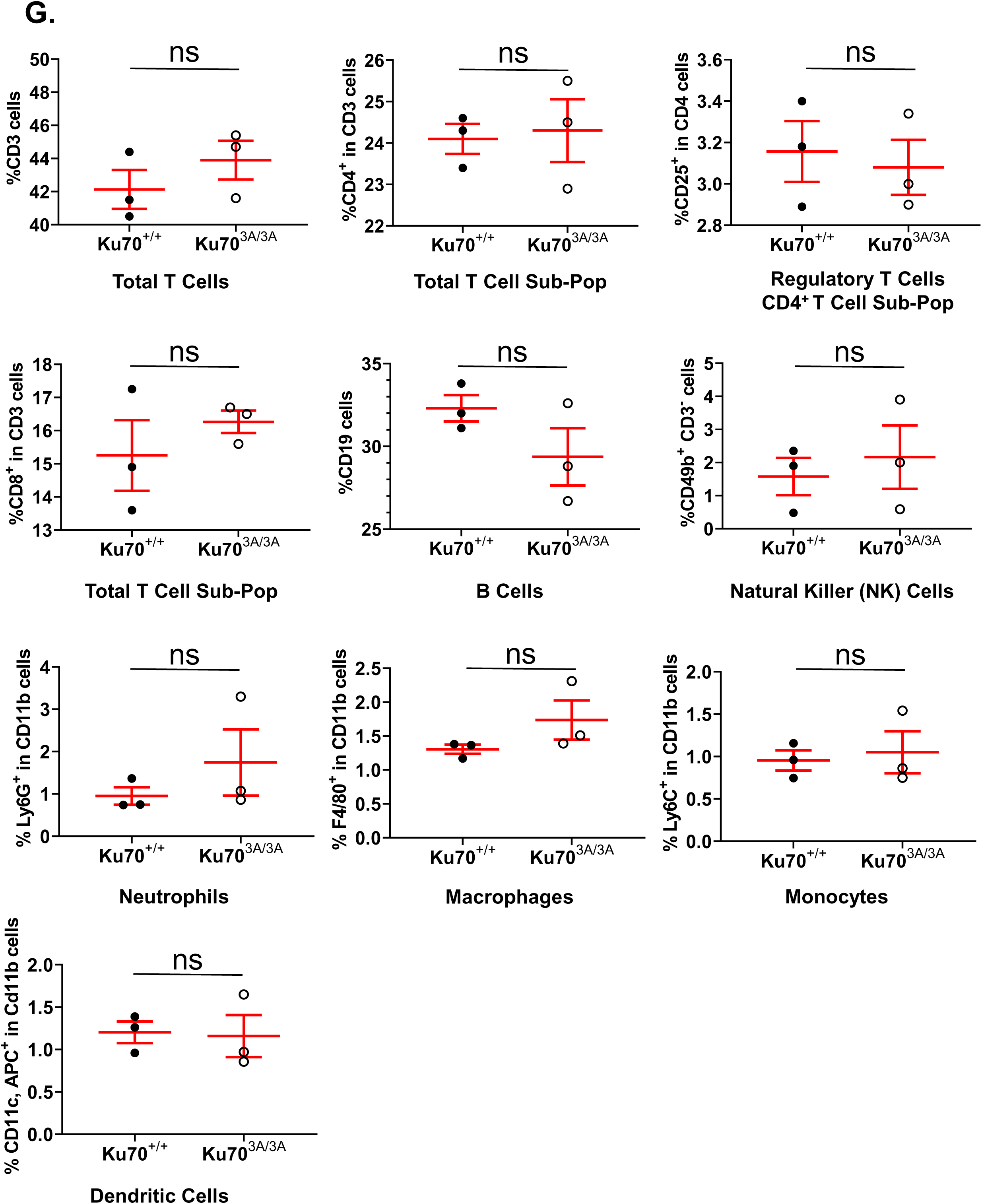
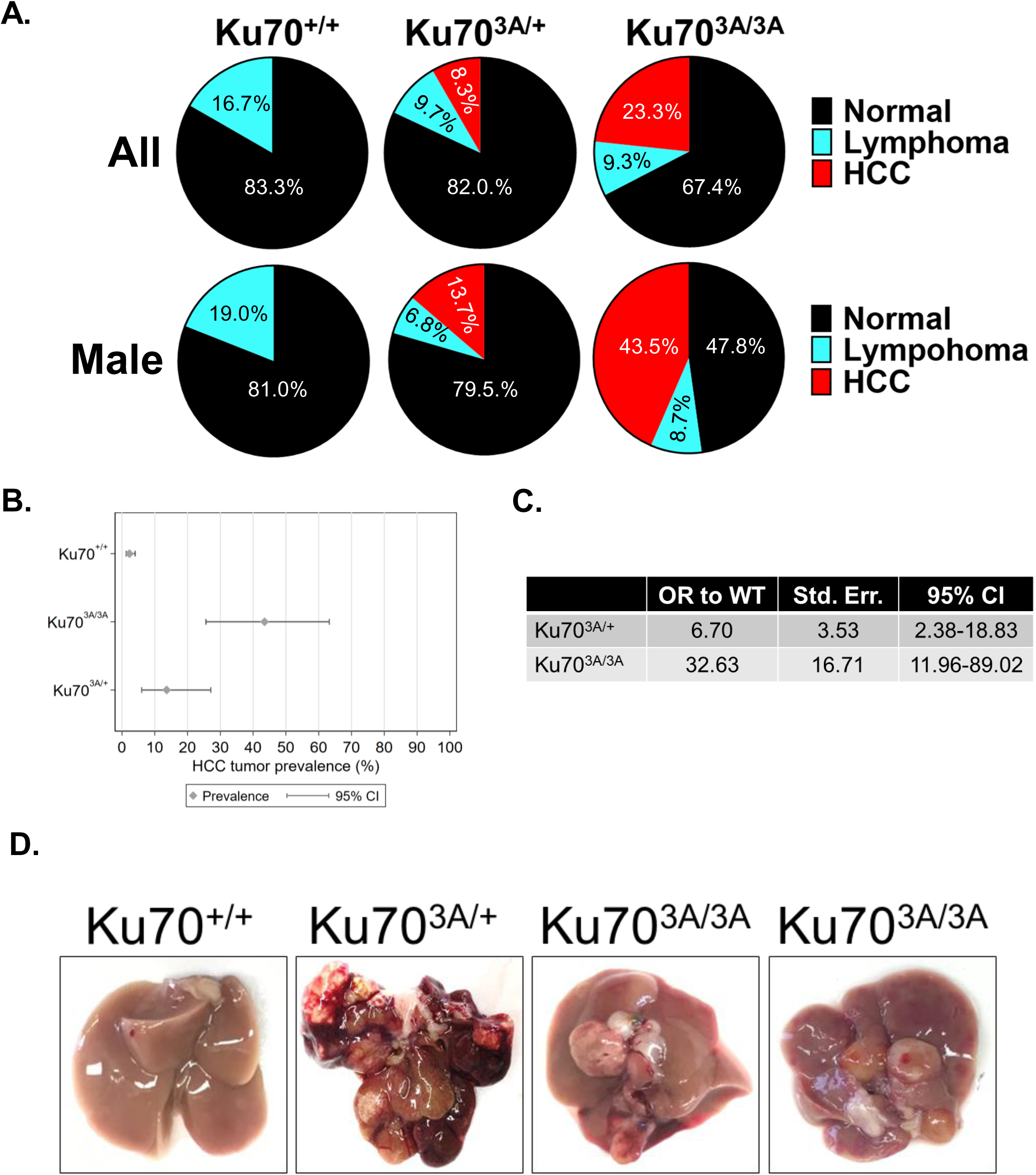
Generation of Ku70^3A/3A^ and Ku70^3A/+^ mice and initial characterization. **A**. Nucleotides changed to ablate phosphorylation of mouse Ku70 phosphorylation sites. Sequencing (**B**) and PCR (**C**) verification of Ku70^3A/3A^ and Ku70^3A/+^ mouse genotypes. **D.** Image of Ku70^+/+^, Ku70^3A/+^, Ku70^3A/3A^ mice. **E.** Growth of male Ku70^+/+^, Ku70^3A/+^, Ku70^3A/3A^ mice was tracked by weight. Error bars denote SD values for the weight of five mice of each genotype. **F.** Kaplan-Meier survival curves for Ku70^+/+^, Ku70^3A/+^, Ku70^3A/3A^ mice for 24 months. **G.** Examination of splenic immune cells from 8-week old Ku70^+/+^and Ku70^3A/3A^ mice. Individual cell populations are marked in each panel. Three mice were used from each mouse genotype and Student t-test was used to examine statistical significance (ns = no statistical difference).

### Lymphocyte development is normal in Ku70^3A/3A^ mice

Since Ku70^-/-^ mice lack mature B cells and serum immunoglobulin and develop only small populations of thymic cells, we next analyzed Ku70^3A/3A^ mice for lymphocyte development (Gu et al., 1997a; Gu et al., 1997b; Li et al., 1998; Li et al., 2007). The spleens of 8-week-old Ku70^+/+^ and Ku70^3A/3A^ mice were isolated and fluorescence activated cell sorting analyses of immune cells were performed. The immune phenotyping analyses of splenic immune cells did not reveal any significant differences in the lymphoid (T-and B-cell populations), T-cell sub-populations (CD4 and CD8 cells; CD4 sub-population CD25 – regulatory T cells) and Natural Killer (NK) cells or myeloid cell populations (neutrophils, macrophages, monocytes and dendritic cells) between Ku70^3A/3A^ and Ku70^+/+^ littermates (Fig. 2G). Together, these results show that the Ku70^3A/3A^ mice do not have a significant defect in V(D)J recombination.

### Ku70^3A/3A^ and Ku70^3A/+^ mice develop spontaneous hepatocellular carcinoma (HCC)

No differences in survival rate was observed in our cohort of mice (Fig. 2F), but necropsy and clinical histopathological analysis surprisingly revealed development of HCC in the Ku70^3A/3A^ and Ku70^3A/+^ mice (**See** Fig. 3D **for representative images**). No HCC was identified in Ku70^+/+^ mice; however, HCC was detected in 8.3% of Ku70^3A/+^ and 23.3% of Ku70^3A/3A^ mice, including HCC being observed as early as 17 months in Ku70^3A/3A^ mice (Fig. 3A). Similar to previous mouse studies, development of HCC occurred more often in male than female mice (Bakiri and Wagner, 2013). Spontaneous HCC was only identified in male mice in the Ku70^3A/3A^ cohort with 43.5% (95% CI for HCC tumor prevalence was 25.6-63.2%) of male mice developing HCC (Fig. 3A). One female in the Ku70^3A/+^ cohort developed HCC and the incidence of male Ku70^3A/+^ developing HCC was 23.3% (95% CI for HCC tumor prevalence was 6.0-27.0%) (Fig. 3A). We performed logistic regression analysis to estimate an odds ratio comparing Ku70^+/+^ to both Ku70^3A/+^ and Ku70^3A/3A^ male mice. As none of our Ku70^+/+^ male mice developed HCC, we used historical data of spontaneous HCC induction in untreated mice in combination with our own data for the logistic regression analysis (Blackwell et al., 1995; Takahashi et al., 2002). The analysis found that the odds of HCC tumorigenesis is 32.63 times larger (95% CI is 11.96-89.02) in Ku70^3A/3A^ mice and 6.7 times larger (95% CI 2.38-18.83) in Ku70^3A/+^ mice compared to the Ku70^+/+^ mice (Fig. 3C). To further characterize the effects of ablation of Ku70 phosphorylation on HCC development, we examined liver sections of Ku70^+/+^ and Ku70^3A/3A^ mice by immunohistochemistry staining with proliferation marker Ki67, DSB marker γH2AX, and free radical damage/reactive oxygen species (ROS) marker 8-oxoguanine (8-oxo-G). Immunohistochemistry analysis of the HCC tumors from Ku70^3A/3A^ revealed strong Ki67, γH2AX, and 8-oxo-G staining, suggesting high levels of proliferation and accumulated DNA damage in the HCC tumors (Fig. 3C). Pathologic analysis also revealed spontaneous lymphoma in all three groups of mice (Fig. 3A), which is not surprising as C57BL/6 mice are prone to lymphomas and hematopoietic neoplasia (Blackwell et al., 1995; Brayton et al., 2012). However, no increase in lymphoma was observed in the Ku70^3A/+^ and Ku70^3A/3A^ compared to the Ku70^+/+^ mice. Moreover, no lymphomas were found in any mouse that was less than 24 months old, suggesting this phenomenon is due to the propensity of aged C57BL/6 to develop lymphoma. Collectively, the data Ku70^3A/3A^ and Ku70^3A/+^ mice develop spontaneous HCC and this correlates with accumulated DNA damage.

**Figure 3.**
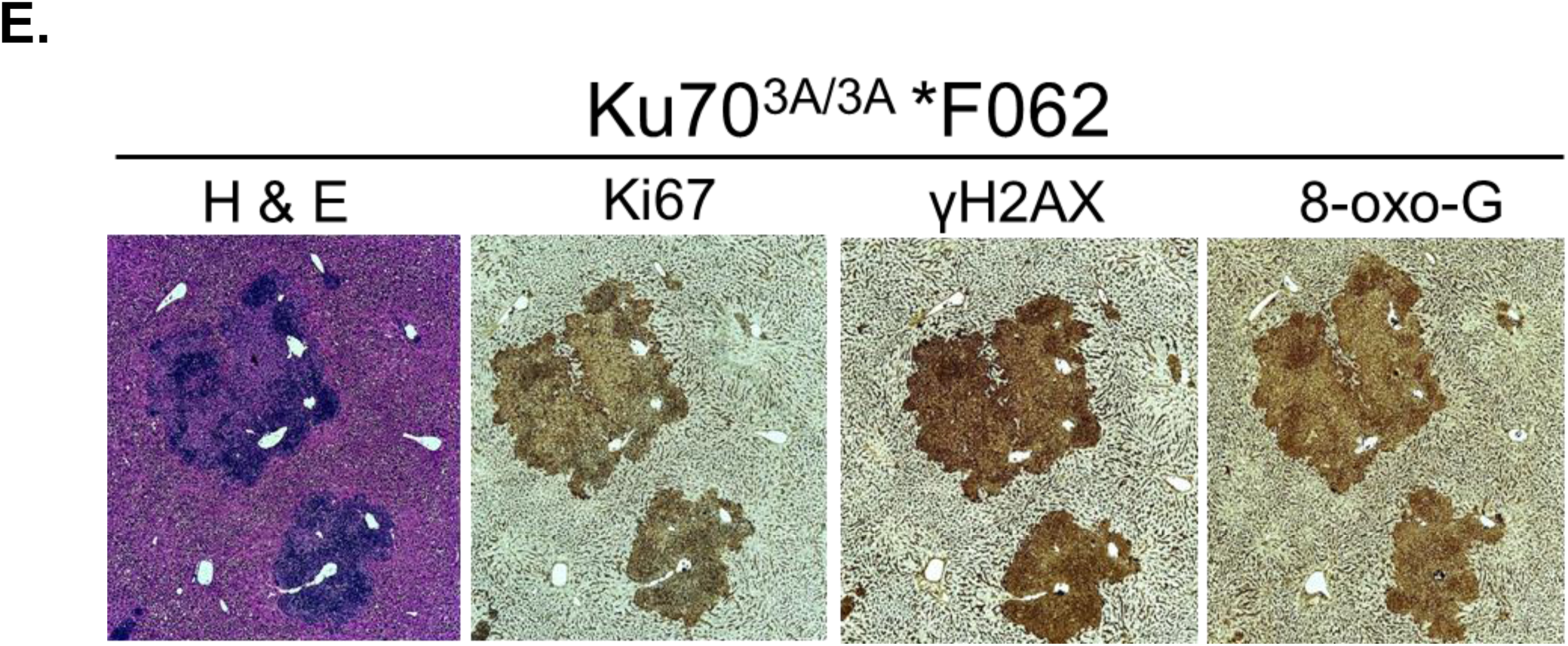
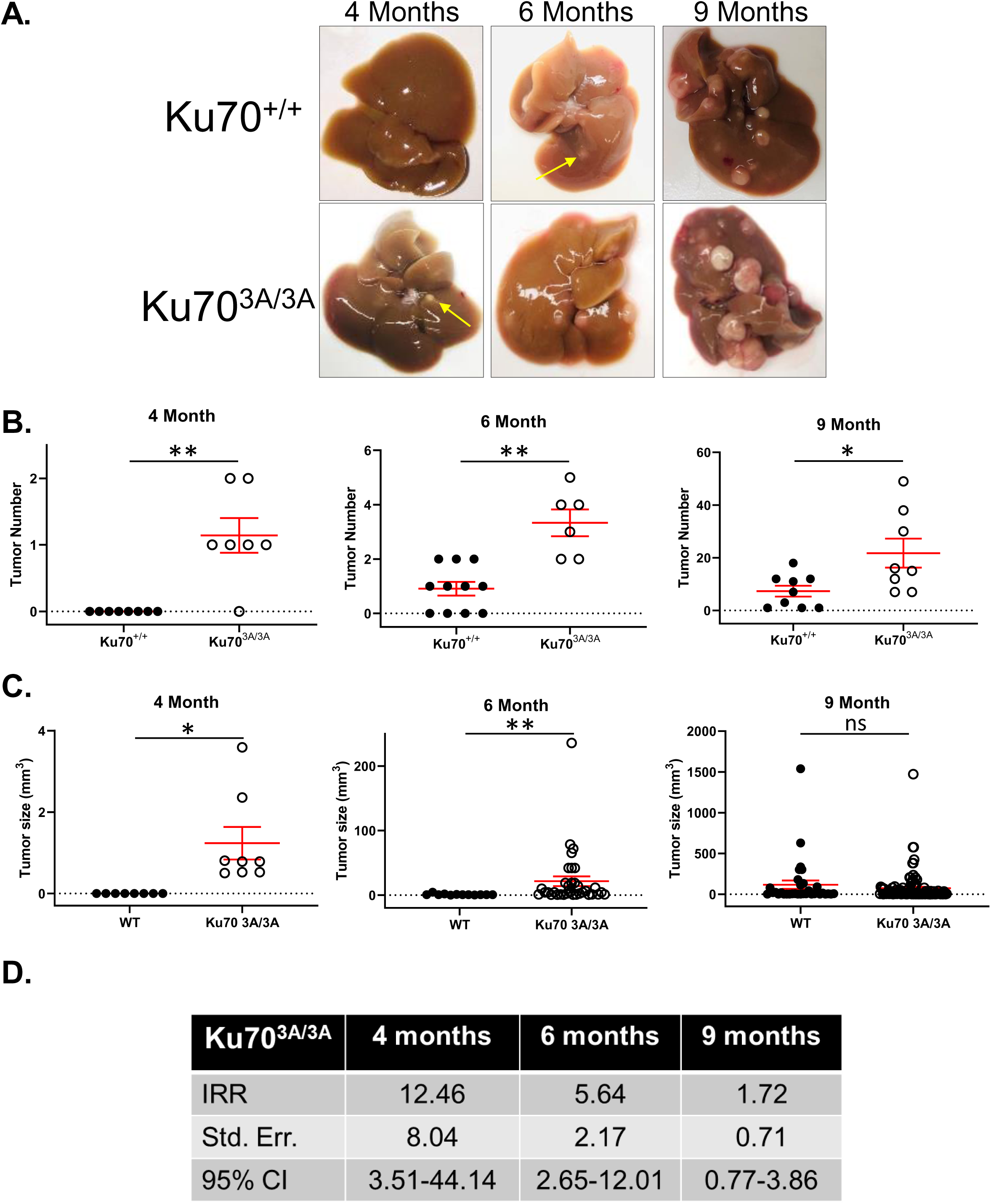
Ku70^3A/3A^ and Ku70^3A/+^ mice have increased incidence of hepatocellular carcinoma (HCC). **A.** Tumor incidence in all (upper panel) and male (lower panel) Ku70^+/+^, Ku70^3A/+^, and Ku70^3A/3A^ mice. **B.** HCC tumor prevalence in Ku70^+/+^, Ku70^3A/+^, and Ku70^3A/3A^ mice with 95% confidence interval (CI). **C.** Logistic regression analysis was performed to examine HCC tumors odds ratio (OR) for Ku70^+/+^, Ku70^3A/+^, and Ku70^3A/3A^ mice. Standard error and 95% CI are also provided. **D.** Representative images of livers and HCCs from Ku70^+/+^, Ku70^3A/+^, and Ku70^3A/3A^ mice. **E.** Immunohistochemistry staining of HCC tumor from Ku70^3A/3A^ F062 mouse. IHC staining was performed using hematoxylin and eosin (H & E) stain and the following antibodies, phospho-Histone H2A.X (Ser139) (γH2AX), Ki67, and 8-oxoguanine (8-oxo-G).

### Ku70^3A/3A^ mice have accelerated chemical-induced HCC development

Since spontaneous HCC development in the Ku70^3A/3A^ mice correlates with accumulated DNA damage and thus defective DNA repair, we next examined if induction of DNA damage via the liver carcinogen DEN promotes accelerated HCC development in Ku70^3A/3A^ mice (Tolba et al., 2015). DEN induces acute liver injury by damaging DNA by directly generating DNA adducts and inter-strand crosslinks (ICLs) and indirectly via ROS production during metabolism of DEN that generates single strand breaks (SSBs) and DSBs (Heindryckx et al., 2009; Santos et al., 2017). DEN works in a dose-dependent manner and DEN-induced HCC development is faster in young mice and males (Heindryckx et al., 2009). 14-day-old Ku70^+/+^ and Ku70^3A/3A^ mice were treated with a single dose (25 mg/kg, intraperitoneally) of DEN and were examined at 4-, 6-, and 9-months post-injection for liver tumors. A minimum of 6 (and up to 11) animals from each genotype were analyzed. There was no evidence of liver neoplasia in the Ku70^+/+^ mice at 4 months, whereas 6 out of 7 Ku70^3A/3A^ mice had at least 1 liver tumor at this time point (average of 1.14 ± 0.26 tumors/mouse) (Fig. 4A **and** B). Furthermore, at 6 months, all (6 out of 6) Ku70^3A/3A^ mice showed liver tumors, with an average of 3.33 ± 0.49 tumors/mouse (Fig. 4A **and** B). At 6 months, 7 out of 11 Ku70^+/+^ mice had liver tumors, but the average was only 0.91 +/-0.25 tumors/mouse. Consistent with an earlier induction of HCC in the mutant mice, tumors found in 6-month-old Ku70^3A/3A^ mice were significantly larger (21.42 ± 7.53 mm^3^) than those found in Ku70^+/+^ mice (0.91 ± 0.34 mm^3^) (Fig. 4C). Expectedly, at 9 months, all mice in each genotype cohort had DEN-induced HCC, but the Ku70^3A/3A^ mice had significantly higher tumor numbers (21.75 ± 5.48 tumors/mouse) compared to Ku70^+/+^ mice (7.33 ± 2.07) with no difference in average tumor size between the two cohorts observed (Fig. 4A-C). Negative binomial regression analysis was utilized to calculate incidence-rate ratio for tumor development at each time point. The incidence rate of DEN-induced tumor formation for Ku70^3A/3A^ mice is 12.5 times (95% CI 3.51-44.14) higher at 4 months, 5.6 times (95% CI 2.65-12.01) higher at 6 months, and 1.7 (95% CI 0.77-3.86) times higher at 9 months compared to Ku70^+/+^ mice (Fig. 4D). Control (DEN-untreated) Ku70^3A/3A^ and Ku70^+/+^ mice did not develop any tumors when the experiment was terminated 9 months post-treatment. Similar to the spontaneous tumors, the DEN-induced HCC in Ku70^3A/3A^ mice showed strong Ki67, γH2AX, and 8-oxo-G staining, showing accumulation of DNA damage (Fig. 4E). Collectively, Ku70^3A/3A^ mice develop spontaneous and have accelerated chemical-induced HCC compared to Ku70^+/+^ littermates.

**Figure 4.**
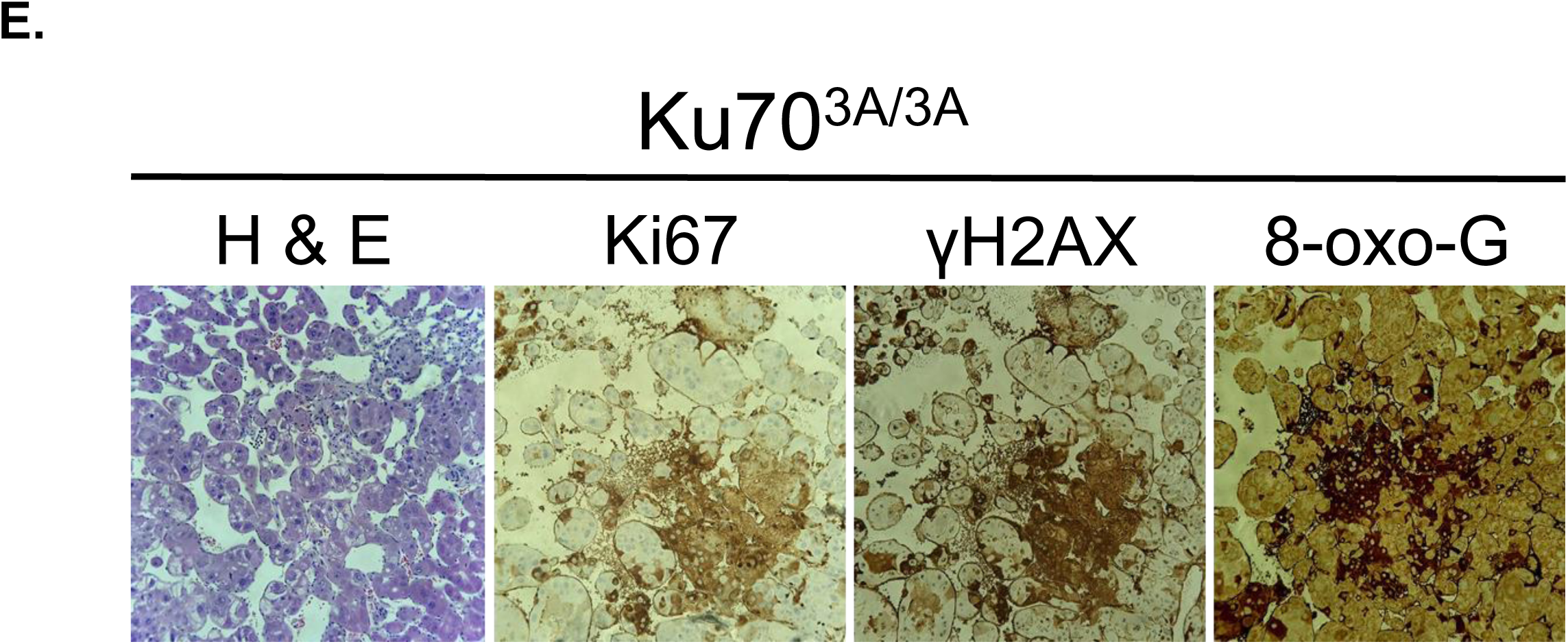
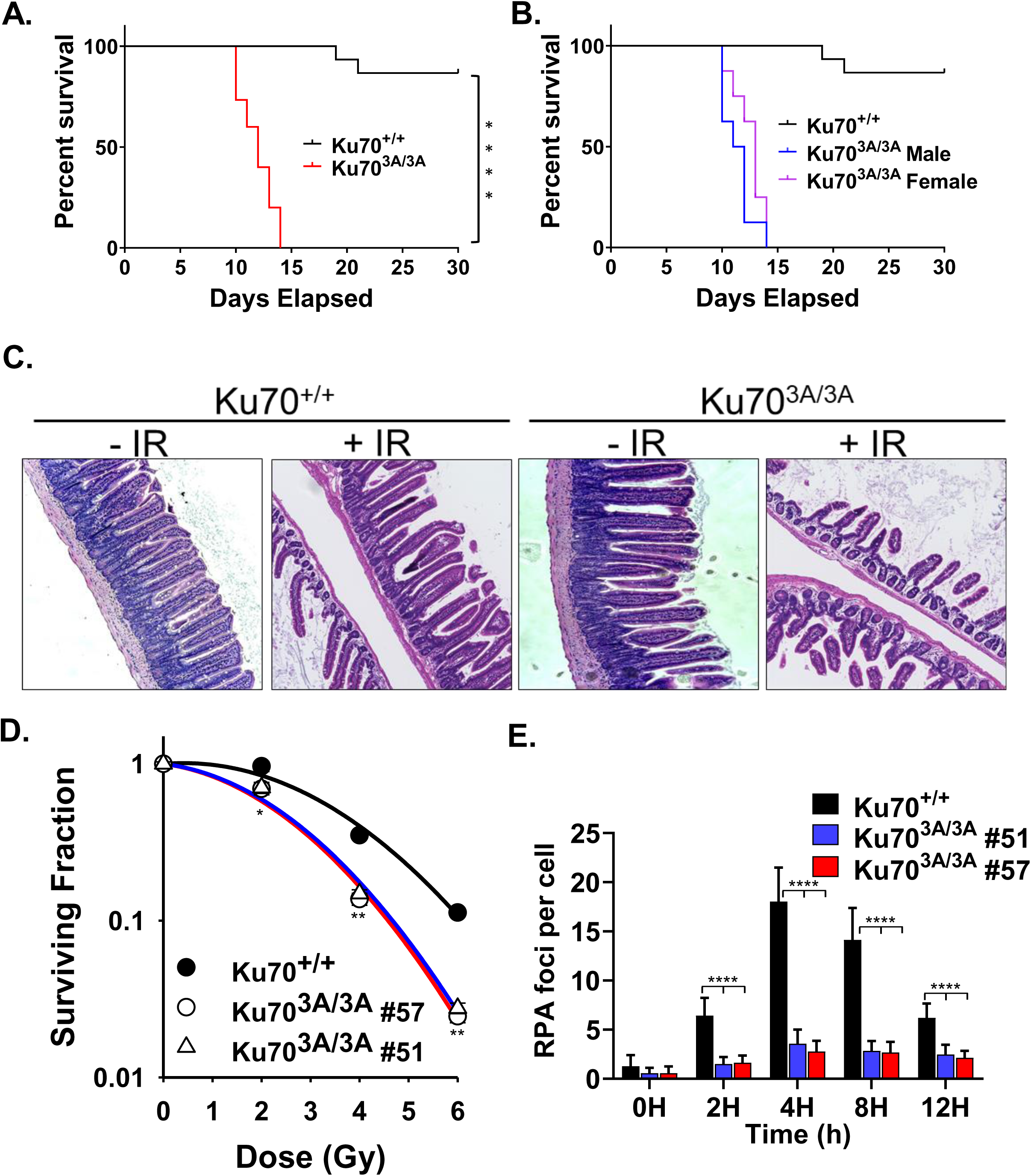
Ku70^3A/3A^ mice have increased accelerated DEN-induced HCC compared to Ku70^+/+^ littermates. **A.** Representative images of livers and HCCs from Ku70^+/+^ and Ku70^3A/3A^ mice 4, 6, and 9 months following injection of the liver carcinogen, DEN. DEN-induced tumor number (**B**) and tumor size (**C**) in Ku70^+/+^ and Ku70^3A/3A^ mice 4, 6, and 9 months following DEN injection. SEM is shown and Welch’s t-test was used to examine statistical significance. (*p < 0.05 and **p < 0.01). **D.** Negative binomial regression was used to calculate incidence rate ratio (IRR) for DEN-induced tumorigenesis. Standard error and 95% CI are also provided. **E.** Immunohistochemistry staining of DEN-induced HCC tumor from Ku70^3A/3A^. IHC staining was performed using hematoxylin and eosin (H & E) stain and the following antibodies, phospho-Histone H2A.X (Ser139) (γH2AX), Ki67, and 8-oxoguanine (8-oxo-G).

### Ku70^3A/3A^ mice are radiosensitive and have attenuated IR-induced DNA end resection

The liver tumors in Ku70^3A/3A^ mice have accumulated DNA damage, suggesting defective DNA repair in this cohort of mice. To test if the Ku70^3A/3A^ mice have defective DSB repair compared to their Ku70^+/+^ littermates, total body irradiation (TBI) experiments were performed. Treatment with 9 Gy of γ-rays revealed that Ku70^3A/3A^ mice are significantly radiosensitive compared to Ku70^+/+^ mice (Fig. 5A). All (15 out of 15) Ku70^3A/3A^ mice succumbed to radiation injury between days 10 and 14, whereas by day 30, only 13.3% (2 out of 15) of Ku70^+/+^ mice had perished from the radiation exposure. No statistical difference between male and female Ku70^3A/3A^ mice in radioresponse was observed (Fig. 5B). The radiosensitivity of the Ku70^3A/3A^ mice correlated with classic acute radiation syndrome (ARS) symptoms, as intestinal H&E staining showed significant intestinal crypt and villi destruction in the Ku70^3A/3A^ mice compared to Ku70^+/+^ (Fig. 5C). Similarly, MEFs isolated from two Ku70^3A/3A^ mice (11.5-day embryo) were radiosensitive compared to those obtained from Ku70^+/+^ littermates (Fig. 5D). Ku70^3A/3A^ MEFs also showed decreased IR-induced RPA foci formation compared to Ku70^+/+^ MEFs, illustrating that blocking Ku70 phosphorylation in the mouse cells results in a significant decrease in DNA end resection (Fig 5E **and. Sup. Fig. 4**). Together, the data illustrates that the Ku70^3A/3A^ mice are radiosensitive and suggests this is due to decreased HR.

**Figure 5.**
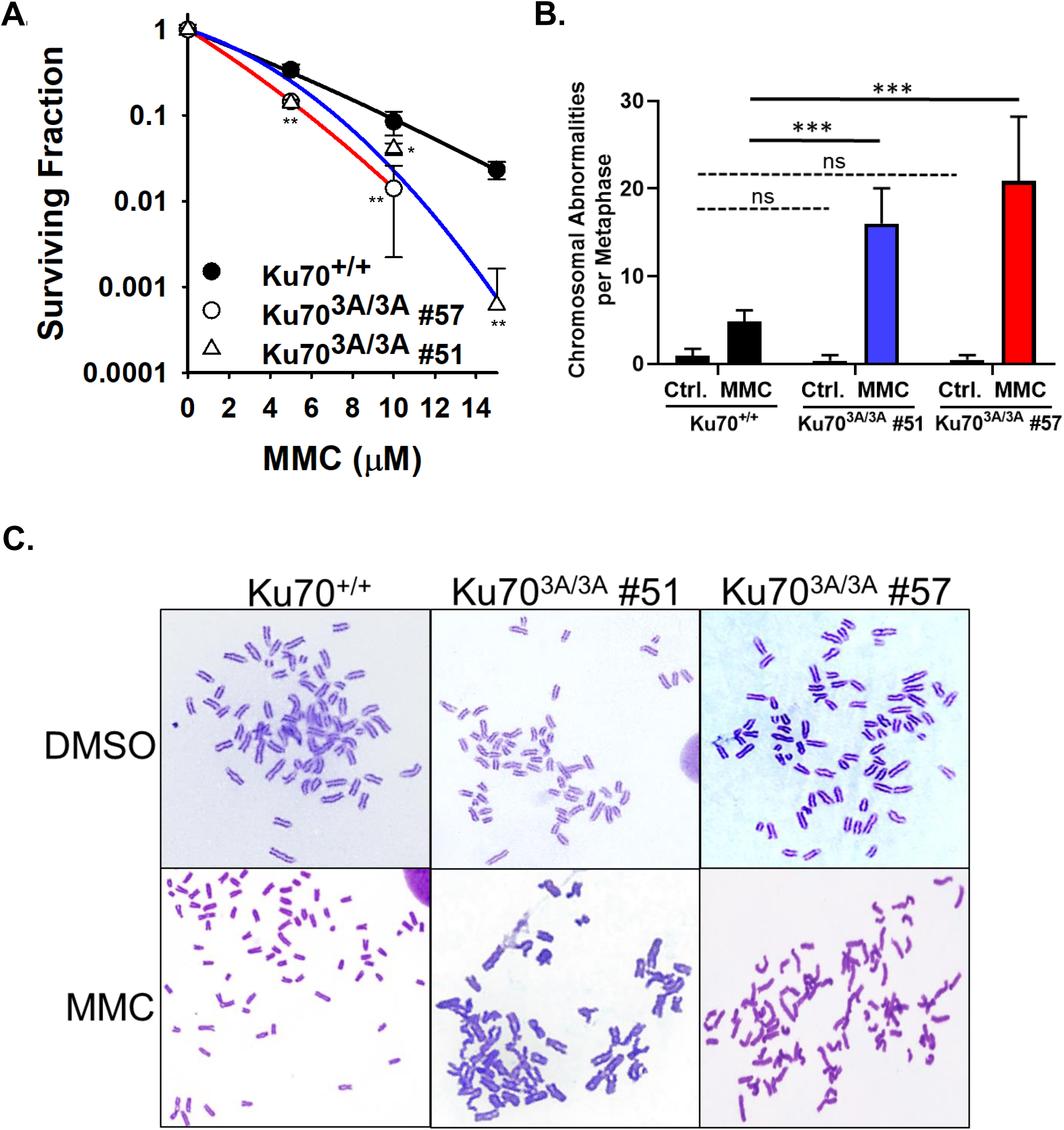
Ku70^3A/3A^ mice are radiosensitive and have attenuated IR-induced DNA end resection. Kaplan-Meier survival curves for all (**A**) and male vs. female (**B**) Ku70^+/+^ and Ku70^3A/3A^ mice following total body irradiation with 9 Gy. Log rank test was used to examine statistical significance. (****p < 0.0001). **C.** Hematoxylin and eosin (H & E) stain of colon of Ku70^+/+^ and Ku70^3A/3A^ mice 10 days following total body irradiation. **D.** Clonogenic survival assays were performed to compare the radiation sensitivities of MEFs isolated from a Ku70^+/+^ mouse or two separate Ku70^3A/3A^ mice (clones #51 and #57). Cells were irradiated at the indicated doses and plated for analysis of survival and colony-forming ability. Error bars denote SD values and Student’s t-test was performed to assess statistical significance (*p < 0.05 and **p < 0.01). (**E**) Immunostaining of RPA foci in Ku70^+/+^ and Ku70^3A/3A^ MEFs after 8 Gy. Cells were pre-extracted and fixed 2, 4, 8, or 12 h after IR and immunostained for RPA in EdU positive cells (S phase). RPA foci were counted for each cell and averaged. Student’s t-test was performed to assess statistical significance (****p < 0.0001). Error bars denote standard error of the mean (SEM) for at least 3 independent experiments.

### Ku70^3A/3A^ mice are defective in homologous recombination (HR) repair

Ku70^3A/3A^ mice are radiosensitive and Ku70^3A/3A^ MEFs have decreased DNA end resection, which we postulate is due to decreased HR. Insufficient transfection rates in the Ku70^3A/3A^ MEFs resulted in the inability to examine HR rates using a HR GFP reporter assay; therefore, we compared the sensitivity of Ku70^3A/3A^ and Ku70^+/+^ MEFs to the DNA cross-linking agent, mitomycin C (MMC). MMC-induced ICLs are effectively resolved/repaired via the Fanconi anemia (FA) pathway in conjunction with HR (Deans and West, 2011). Thus, an increase in MMC sensitivity can be used as an indirect test for HR-deficiency. As shown in Fig. 6A, the two Ku70^3A/3A^ MEF cell lines are more sensitive to MMC than Ku70^+/+^ MEFs. Furthermore, the Ku70^3A/3A^ MEFs showed significantly higher MMC-induced chromosomal abnormalities compared to their Ku70^+/+^ counterparts (Fig 6B-C). The Ku70^+/+^ MEFs had 4.85 ± 1.26 chromosomal abnormalities after MMC treatment, whereas Ku70^3A/3A^ #51 and #57 had 16.00 ± 4.00 and 20.84 ± 7.32 chromosomal abnormalities, respectively. Together, our data illustrates that blocking Ku70 phosphorylation in a mouse model results in decreased DNA repair capacity, which induces chronic DNA damage and increased genomic instability driving the induction of HCC.

**Figure 6.**
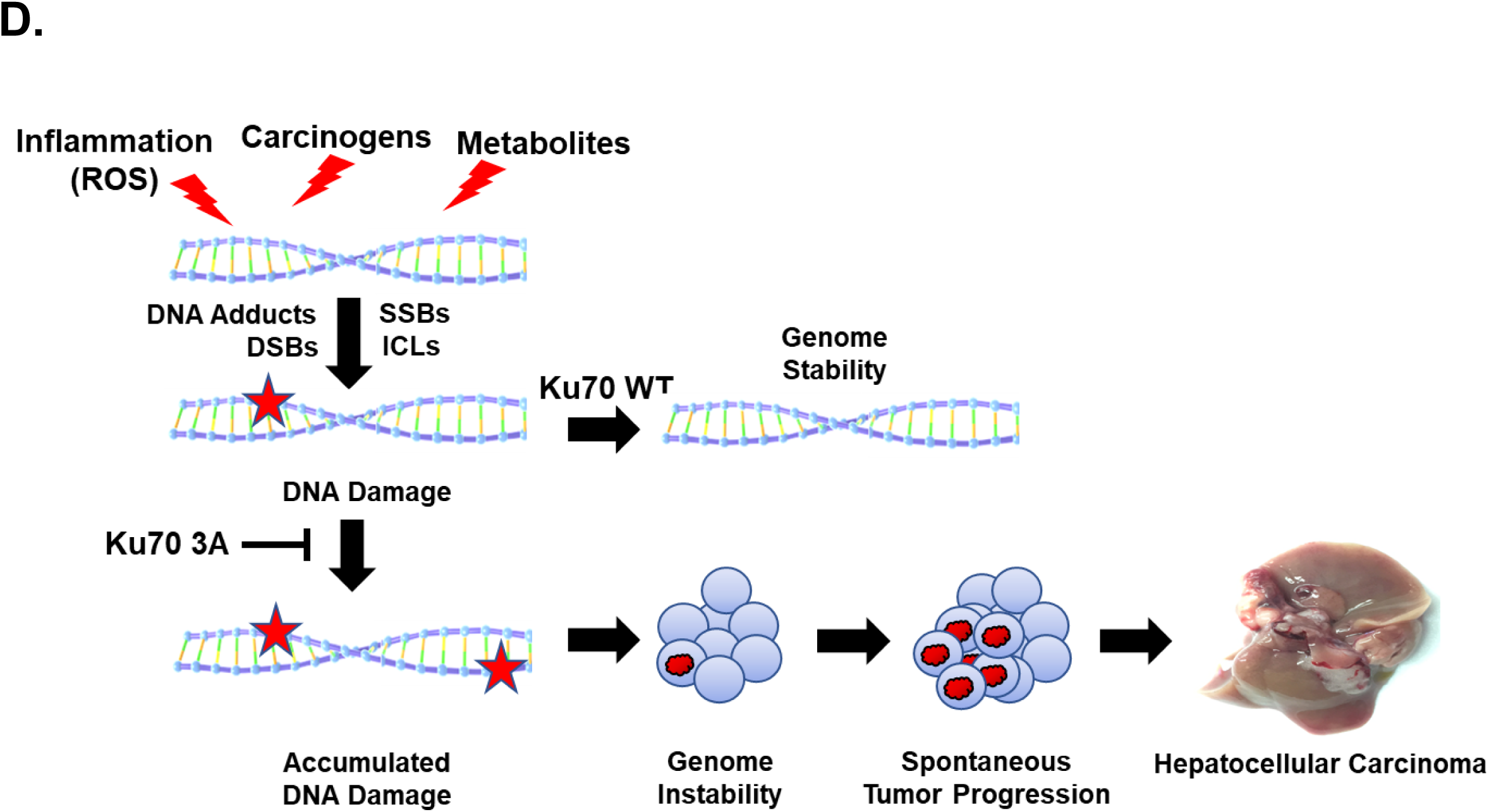
Ku70^3A/3A^ mice are sensitive to the DNA cross-linking agent mitomycin C (MMC). **A.** Clonogenic survival assays were performed to compare the radiation sensitivities of MEFs isolated from a Ku70^+/+^ mouse or two separate Ku70^3A/3A^ mice (clones #51 and #57). Cells were treated with MMC at the indicated doses and plated for analysis of survival and colony-forming ability. Error bars denote SD values and Student’s t-test was performed to assess statistical significance (*p < 0.05 and **p < 0.01). **B.** Metaphase spreads were performed following treatment with MMC in Ku70^+/+^ and Ku70^3A/3A^ MEFs and chromosomal abnormalities were enumerated. Error bars denote SEM values and Welch’s t-test was performed to assess statistical significance (no = no significance, ***p < 0.001). **C.** Representative images of chromosomes following treatment with mock (DMSO) or MMC. **D.** Model.

## DISCUSSION

Liver cancer is the fifth leading cause of cancer-related deaths (Llovet et al., 2016). Among primary liver cancers, HCC is the most common subtype and is associated with multiple risk factors, including hepatitis B or C infection, alcohol consumption, gender (male), metabolic disorders (obesity and diabetes), and diet contamination (aflatoxins) (Llovet et al., 2016). The major HCC risk factors damage DNA either directly, such as DNA alkylating agents, or indirectly via inflammation, which promotes the release of DNA damaging ROS. The promotion of DNA damage and genomic instability are closely associated with HCC development (Rao et al., 2017). However, the factors required for protecting the liver from DNA damage and thus suppress HCC development and progression are mostly unknown. A characteristic trait in genetic alterations in liver cancer is the broad spectrum of mutations. The majority of the ‘most frequently mutated genes’ occur in <10% of HCCs, suggesting there is not one primary driver of liver carcinogenesis but it due to liver injury via chronic inflammation and DNA damage (Rao et al., 2017).

In this study, we developed a mouse model using C57BL/6 mice in which we blocked phosphorylation of the three conserved putative Ku70 phosphorylation sites by mutating the sites to alanine. The goal was to examine if blocking Ku70 phosphorylation attenuated HR, resulting in genomic instability and carcinogenesis. The Ku70^3A/3A^ mice are distinct from Ku70^-/-^ (null) mice, as blocking Ku70 phosphorylation does not drive gross defects in immune cell production or premature aging, suggesting that V(D)J recombination and telomere maintenance are not affected, respectively (Gu et al., 1997b; Li et al., 1998; Li et al., 2007). However, Ku70^3A/3A^ and Ku70^3A/+^ developed spontaneous HCC and Ku70^3A/3A^ mice had accelerated DEN-induced HCC compared to Ku70^+/+^ mice. Liver tumors are extremely rare in untreated C57BL/6 male mice; the incidence of liver tumors by 2 years of age is <5%, and DEN-induced HCC is also lower in C57BL/6 compared to other strains (Blackwell et al., 1995; Brayton et al., 2012; Drinkwater and Ginsler, 1986; Takahashi et al., 2002). Since the C57BL/6 mouse line is resistant to liver carcinogenesis, our data indicate that the Ku70 3A mutations specifically drives HCC development and is not due to intrinsic increased susceptibility of the mouse strain. Logistic regression analysis found spontaneous HCC tumor odds to be 33.6 times larger in Ku70^3A/3A^ compared to Ku70^+/+^ mice, showing the significance of the Ku70 3A mutations in promoting HCC.

Mouse models have allowed groups to study the physiological functions of DSB repair proteins and to characterize their roles in protecting against cancer (Specks et al., 2015). The majority of these studies have utilized knockout of the perspective gene, with limited studies using mouse models with point mutations in DSB repair proteins. Studies using mouse models examining the role of the DDR machinery in modulating liver carcinogenesis are limited. DEN-induced hepatocarcinogenesis is abrogated in ATM-deficient mice (Teoh et al., 2010). Ku70^-/-^, Ku80^-/-^, and Ku70^-/-^Ku80^-/-^ mice with the same genetic background show that loss of Ku promotes early aging in each cohort of mice, but without substantially increased cancer levels (Li et al., 2007). It should be noted, in the Ku70^-/-^ cohort, one mouse developed HCC at 40 weeks (Li et al., 2007). Ku80^+/-^PARP^-/-^ mice develop liver cancers, including HCC (Tong et al., 2002). Furthermore, Ku70^-/-^ mice had accelerated DEN-induced HCC development compared to Ku70^+/+^ and Ku70^+/-^ littermates (Teoh et al., 2008). These reports indicate that Ku may play an important role in protecting against HCC tumorigenesis. Although point mutations in many DSB repair proteins are believed to drive tumorigenesis in humans, surprisingly, few studies to date have found point mutations in DDR genes that result in spontaneous tumorigenesis in mice. Ablating the CHK2 phosphorylation site at S971 in *BRCA1* resulted in increased uterus hyperplasia and ovarian abnormalities in mice, and accelerated tumorigenesis following treatment with IR compared to wild-type littermates (Kim et al., 2004). The *BRCA1^C61G^* mutation in the RING domain of BRCA1 is a common pathogenic missense mutation in familial breast cancer (Krais and Johnson, 2020). Introduction of this mutation in mice resulted in embryonic lethality, and a conditional *BRCA1^C61G^* mutation in mammary cells in a p53 null background resulted in spontaneous mammary tumors (Drost et al., 2011). Congenital bone marrow failure and rapid death was observed in mice in which DNA-PK_cs_ phosphorylation at the T2609 cluster was ablated and rescuing these mice with a bone marrow transplant to prevent early mortality resulted in an increase in spontaneous tumorigenesis (Zhang et al., 2016a; Zhang et al., 2011). To our knowledge, this present study is the first to report spontaneous induction of HCC due to point mutations in a DDR protein. Lastly, the data presented in this manuscript shows that blocking Ku70 phosphorylation results in increased spontaneous and DEN-induced HCC, which together with the data in the literature, suggest that the Ku heterodimer plays a key function in suppressing HCC and strengthens its perceived role as a tumor suppressor.

The role of Ku in the pathogenesis and development of human HCC remains unclear. One report found that Ku80 is frequently downregulated in human HCC and this downregulation significantly correlated with HBV and liver cirrhosis (Wei et al., 2012). However, a different study found that Ku70 upregulation correlated with HCC and was associated with poor prognosis in HCC patients (Zhang et al., 2016b). Differential expression of the canonical NHEJ factors, Ku70, Ku80, DNA-PK_cs_, and XRCC4, correlates with increased susceptibility to HCC and liver carcinogens, and increased unrepaired DNA damage such as DSBs and DNA adducts (Sishc and Davis, 2017). Sequencing of tumor samples has been extremely beneficial in identifying mutations in DDR genes that are potential drivers of carcinogenesis, including HCC. Data mining of the COSMIC (Catalogue of Somatic Mutations in Cancer) and TCGA databases identified multiple single nucleotide variations in the *XRCC6* (Ku70) gene in and near the Ku70 phosphorylation cluster, including at the conserved phosphorylation sites S314 (endometrial cancer) and T316 (colorectal cancer) (**Sup. Fig. 5**). Mutations at G309 and S324 were identified in patients with HCC (**Sup. Fig. 5**). Collectively, this supports the notion that Ku70 phosphorylation sites and amino acid near these sites are important for protecting against carcinogenesis. Unfortunately, many targeted sequencing programs do not include the canonical NHEJ factors in their “key cancer genes”, including the Memorial Sloan Kettering-Integrated Mutation Profiling of Actionable Cancer Targets (MSK-IMPACT) (Cheng et al., 2015). Similarly, a clinical trial found that alterations in genes required for BER, FA, HR, and ATM correlate with increased HCC, increased tumor mutation burden, and poorer survival, but again, the canonical NHEJ factors were not included in the study (Lin et al., 2019). As there is interplay between the DNA repair pathways and the fact that mounting evidence, including this study, suggests the DNA-PK complex plays a role in protecting against tumorigenesis and cancer progression, we recommend adding the canonical NHEJ factors, in particular *XRCC6* (Ku70), *XRCC5* (Ku80), and *PRKDC* (DNA-PK_cs_), to the sequencing studies examining mutations of DDR proteins in carcinogenesis (Hsu et al., 2012; Sishc and Davis, 2017).

We propose the following model for the role that Ku70 phosphorylation plays in protecting cells from HCC (Fig. 6D). The liver is constantly under attack from multiple agents, such as carcinogens (aflatoxins), metabolites (aldehydes from alcohol metabolism), and reactive oxygen species (ROS) from inflammation due to obesity. We postulate that these agents generate various DNA damage including DNA adducts, SSBs, DSBs, and ICLs. In Ku70^+/+^ mice, the DNA damage is rapidly repaired and genomic stability is maintained in the liver. However, in Ku70^3A/3A^ mice, the Ku70 phosphorylation mutant attenuates the repair of DSBs that are directly generated or those created due to the processing of DNA adducts, SSBs, and or ICLs, resulting in accumulated DNA damage. This is supported by our data showing that the HCC tumors identified in Ku70^3A/3A^ mice have increased DSBs as assessed by γH2AX staining. Ku70^3A/3A^ mice and MEFs show sensitivity to agents that generate multiple types of DNA damage including DEN (DNA adducts, DSBs, SSBs, and ICLs), IR (SSBs, DSBs, and base damage), and MMC (ICLs and ROS-induced DNA damage), which further supports that Ku70^3A/3A^ mice have issues repairing multiple types of DNA damage and this is likely due to attenuation of HR (Lee et al., 2016). One of the main drivers of repair of MMC-induced DNA damage is the FA pathway, which is responsible for recognition and excision of the DNA crosslinks generated by MMC and the repair is completed by HR (Deans and West, 2011). Patients with FA have a predisposition to liver tumors and *FANCD2^-/-^* mice develop hepatic adenoma and HCC, supporting a role that disrupting the FA pathway and thus the repair of ICLs promotes HCC induction (Alter, 2003; Houghtaling et al., 2003; Kutler et al., 2003; Masserot-Lureau et al., 2012; Rosenberg et al., 2003). Acetaldehyde, an endogenous and alcohol-derived metabolite, generates DNA damage including DNA crosslinks and DNA adducts that when processed can produce DSBs (Wang et al., 2000). In mice, Ku70 cooperates with the FA pathway in mediating the cellular resistance to endogenous and acetaldehyde-induced DNA damage (Garaycoechea et al., 2018). It is also possible that Ku70 3A directly regulates BER, as well as DSB repair pathways. In a p53-mutant background, Ku70^-/-^ mice exhibited an elevated level of point mutations, suggesting a defect in BER, and increased chromosomal rearrangements in liver, but not brain (Choi et al., 2014). The Ku heterodimer has 5’dRP/AP lyase activity and is essential for efficient removal of AP sites near double-strand breaks (Roberts et al., 2010). *Ku80^−/−^* MEFs are hypersensitive to a variety of agents that generate base lesions and SSBs including ROS-inducing agents H_2_O_2_, streptonigrin, and paraquat, and the alkylating agents (methyl methane sulfonate, MMS) and *N*-ethyl-*N*-nitrosourea (ENU) (Li et al., 2013). Ultimately, the attenuation of various DNA damages by the Ku70 3A protein results in accumulated DNA damage, genome instability, spontaneous neoplasia, and finally hepatocellular carcinoma. This study thus identifies a novel role for Ku70 in protecting against HCC.

## CONTRIBUTION OF AUTHORS

A.J.D. and J.S. conceived the study. J.S., J.B., S-Y.W., and A.J.D. conducted the experiments, and collected and analyzed the data. L.J.C. performed the statistical analysis. P.G. performed the clinical histopathological analysis. A.J.D. wrote the manuscript.

## ACKNOWLEDGEMENTS

We thank Drs. Faya Zhang and Raquibul Hannan from the Department of Radiation Oncology at UTSW with sharing their expertise with lymphocyte isolation and staining experiments. We acknowledge the assistance of the UTSW Tissue Management Shared Resource, a shared resource at the Simmons Comprehensive Cancer Center, which is supported in part by the National Cancer Institute under award number P30 CA142543. We also thank Dr. Mat Augustine in the Department of Surgery at UTSW for comments on the manuscript. Finally, we are indebted to Drs. David J. Chen and Benjamin P.C. Chen for their mentorship, guidance on the initiation of the project, and discussions. This research was supported by grants from the National Institutes of Health CA092584, CA162804, and GM04725 to A.J.D.

