## Supplemental Data for "Ablating Ku70 phosphorylation results in defective DNA damage repair and spontaneous induction of hepatocellular carcinoma"

#### SUPPLEMENTARY FIGURE LEGENDS

**Supplementary Figure 1.** Alignment of Ku70 DNA sequences across multiple organisms. Phosphorylation sites are in red.

**Supplementary Figure 2.** Panel of images depicting RPA (A) and Rad51(B) foci in Ku70$^{-/-}$ MEFs or Ku70$^{-/-}$ MEFs complemented with Ku70 wild-type (Ku70$^{+/+}$) or 3A (Ku70$^{3A/3A}$) in EdU positive cells after 8 h after exposure to 8 Gy.

**Supplementary Figure 3.** Growth of female Ku70$^{+/+}$, Ku70$^{3A/+}$, Ku70$^{3A/3A}$ mice was tracked by weight. Error bars denote SD values for the weight of five mice of each genotype.

**Supplementary Figure 4.** Panel of images depicting RPA foci in in Ku70$^{+/+}$ and Ku70$^{3A/3A}$ MEFs after 8 Gy in EdU positive cells after 8 hr.

**Supplementary Figure 5.** Data mining of the COSMIC (Catalogue of Somatic Mutations in Cancer) and TCGA databases in order to identify single nucleotide variations in the *XRCC6* (Ku70) gene in and near the Ku70 phosphorylation cluster,

### Supplementary Figure 1.

| | | |
|---|---|---|
| | 305, 306, 307, 314, 316 | |
| ***H. sapien*** | 296-VKTKTRTFNTSTGGLLLPSDTKRSQIYGSRQ-326- | human |
| ***P. troglodyes*** | 296-VKTKTRTFNTSTGGLLLPSDTKRSQIYGSRQ-326- | chimpanzee |
| ***G. gorilla*** | 296-VKTKTRTFNTSTGGLLLPSDTKRSQIYGSRQ-326- | gorilla |
| ***D. leucas*** | 294-VKTKTRTFNVNTGSLLLPSDTKRSQTYGSRQ-324- | beluga whale |
| ***S. scrofa*** | 292-VKTKTRTFNVNTGSLLLPSDTKRSQTYGNRQ-322- | pig |
| ***R. norvegicus*** | 294-VKTKTRTFNVNTGSLLLPSDTRSLTGTRQIV-324- | rat |
| ***M. musculus*** | 294-VKTKTRTFNVNTGSLLLPSDTKRSLTYGTRQ-324- | mouse |
| ***G. gallus*** | 333-VKTKTRVFNGKTGSLLLPSDTKRAQTYGNRG-363- | chicken |
| ***V. komodoensis*** | 294-VKTKTRTFSRETGGLLLPSDTKRAQIYGNRQ-324- | komodo dragon |
| ***O. tshawytscha*** | 292-VRTKTRLYHTQTGSLLLPSDTKRVQVYAGRQ-322- | salmon |
| ***E. lucius*** | 292-VRTKTRLYHTQTGSLLLPSDTKRAQVYASKQ-322- | northern pike |
| ***D. rerio*** | 293-VRTKSRLFHTQTGGILLPNDTKRAQVYGQKQ-323- | zebrafish |
| ***X. laevis*** | 294-VKTKTRIFHSNTGSLLLPSDTKRSQTYGNRQ-324- | horned frog |
| ***D. melanogaster*** | 295-VRTKRVITVQDDGSQDIETGGWYTCNVGERD-335- | fruit fly |
| ***C. elegans*** | 265-KIVKTSGYVKLEDSIRNRRDLKKSIEIGGEK-345- | worm |
| ***A. thaliana*** | 307-VKVERS-YICTDTGAIMQDPIQRIQPYKNQN-336- | arabidopsis |
| ***S. cerevisiae*** | 311-EAYSKRKFLNPITGEDVTGKTVKVVPYGDLD-341- | yeast |

### Supplementary Figure 2.

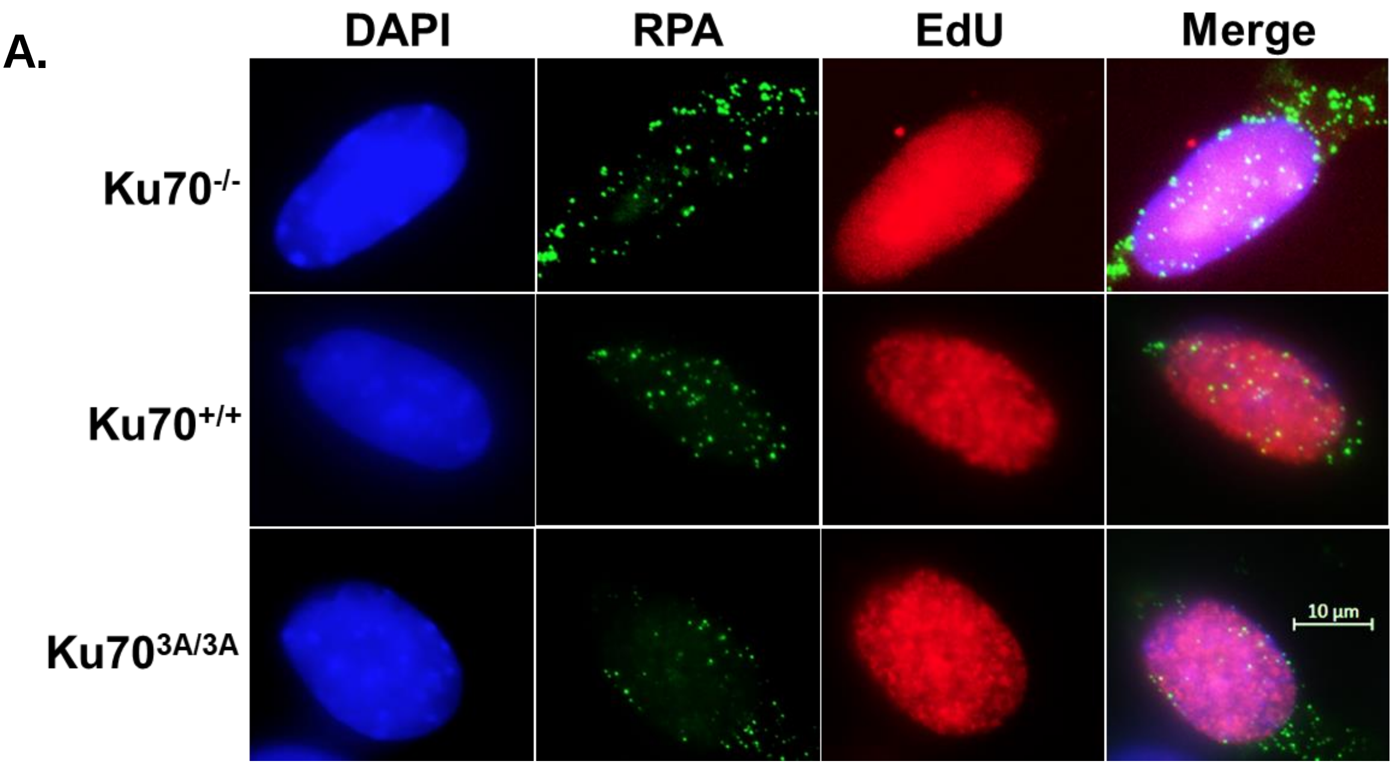


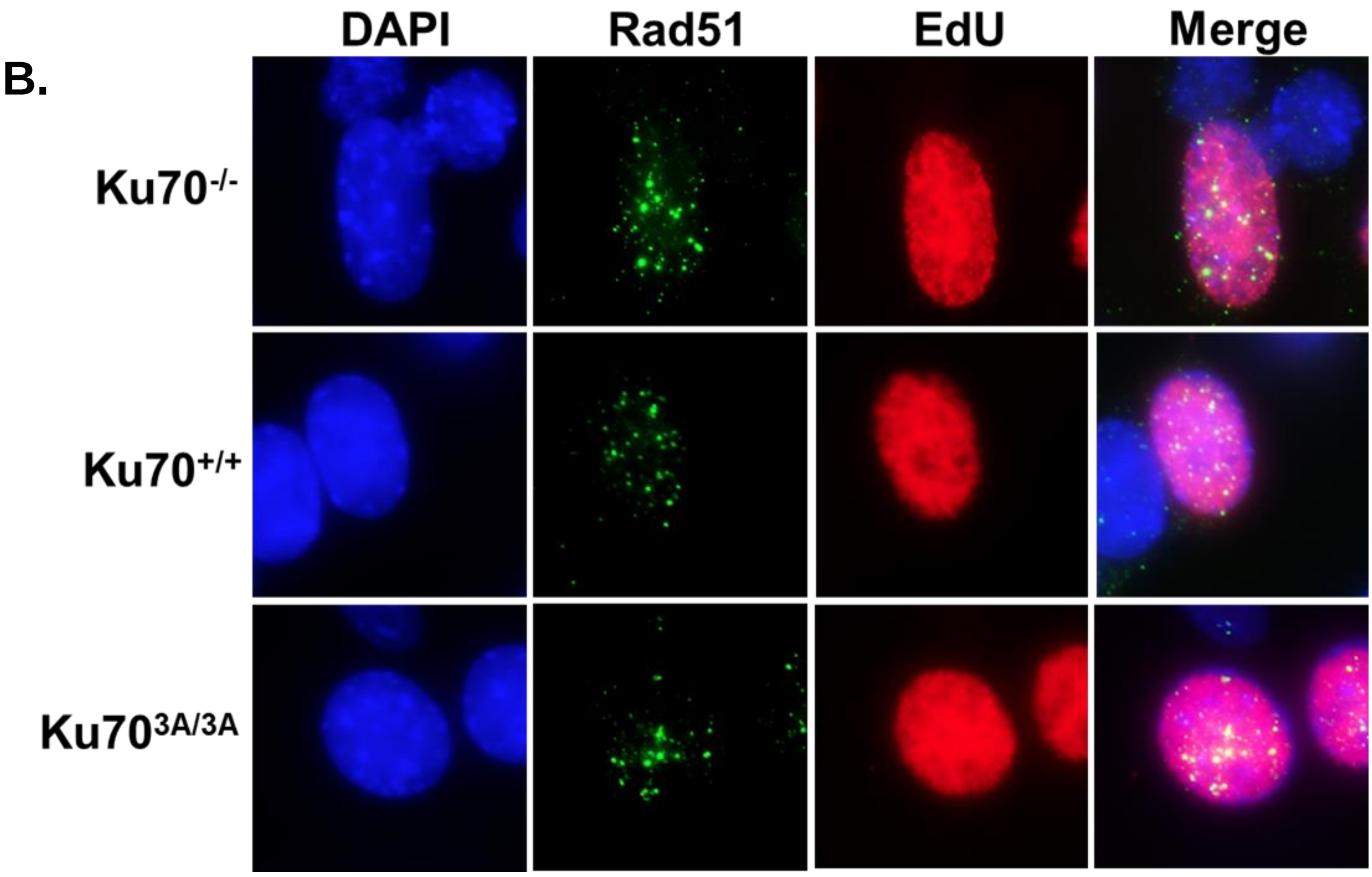

#### Supplementary Figure 3.

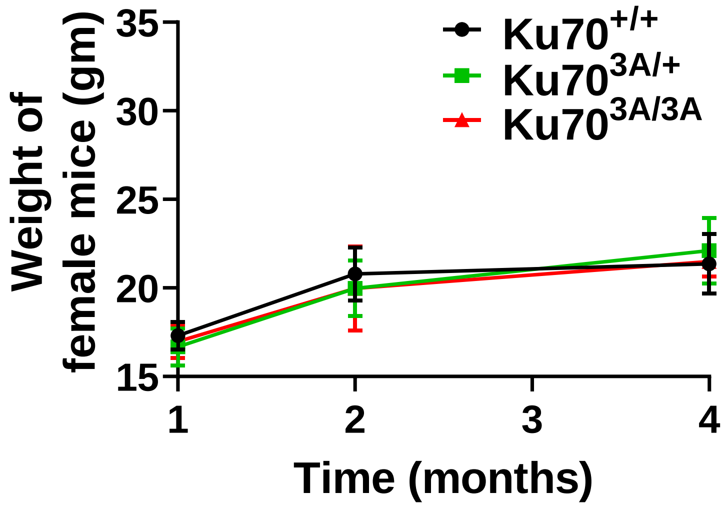

### Supplementary Figure 4.

A.

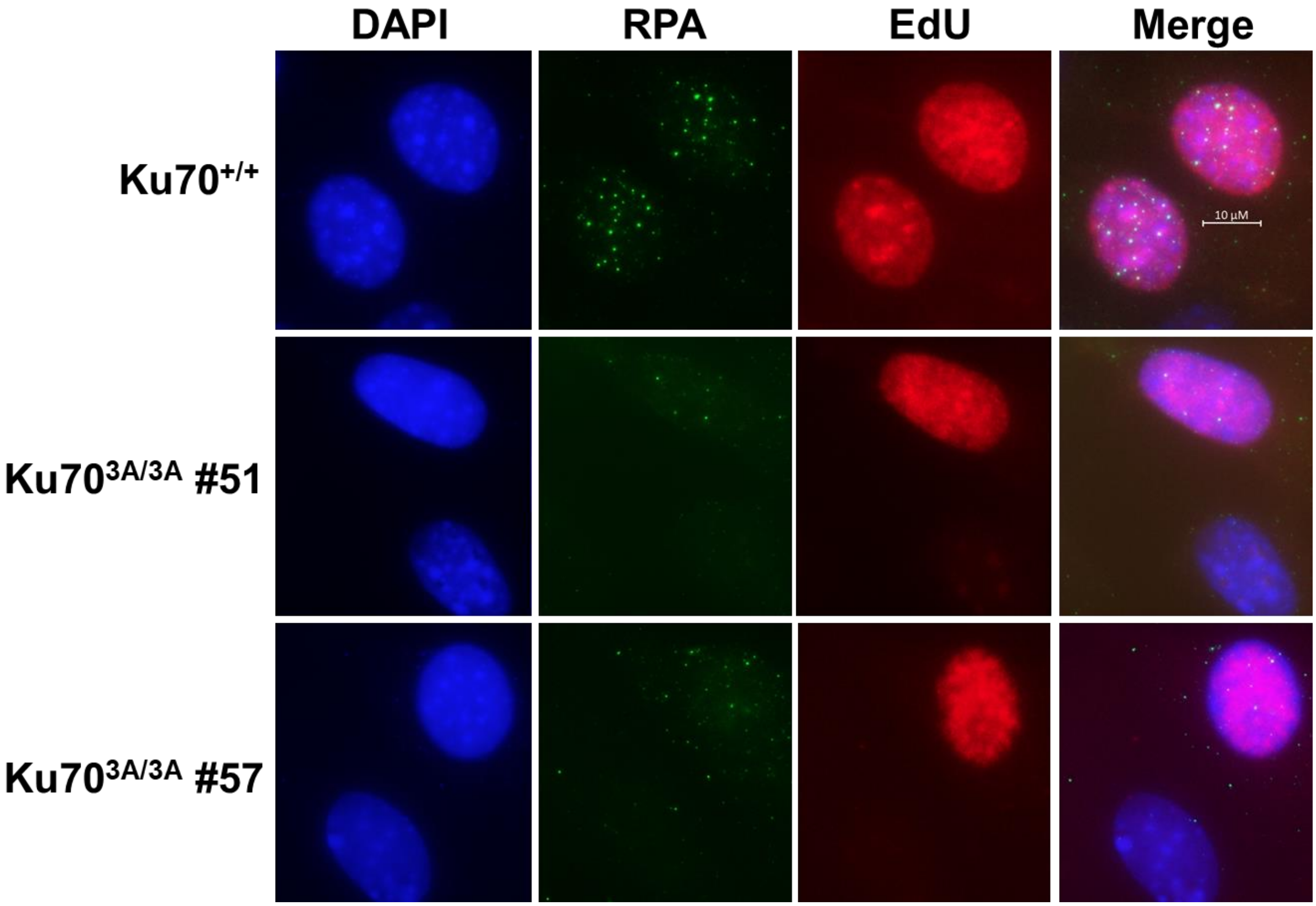

#### Supplementary Figure 5.

306 305 307 314 316

296-VKTKTRTFNTSTGGLLLPSDTKRSQIYGSRQ-326

| Mutation | Cancer Type |
|---|---|
| V296M | Melanoma |
| T302I | Ovarian |
| G309C | Liver |
| S314R | Endometrial |
| D315N | Glioblastoma |
| T316I | Colorectal |
| K317N | Myeloma |
| S324N | Liver |
| R325C | Colorectal |
